# Skin-associated *Corynebacterium amycolatum* shares cobamides

**DOI:** 10.1101/2024.04.28.591522

**Authors:** MH Swaney, N Henriquez, T Campbell, J Handelsman, LR Kalan

## Abstract

The underlying interactions that occur to maintain skin microbiome composition, function, and overall skin health are largely unknown. Often, these types of interactions are mediated by microbial metabolites. Cobamides, the vitamin B_12_ family of cofactors, are essential for metabolism in many bacteria, but are only synthesized by a small fraction of prokaryotes, including certain skin-associated species. Therefore, we hypothesize that cobamide sharing mediates skin community dynamics. Preliminary work predicts that several skin-associated *Corynebacterium* species encode *de novo* cobamide biosynthesis and that their abundance is associated with skin microbiome diversity. Here, we show that commensal *Corynebacterium amycolatum* produces cobamides and that this synthesis can be tuned by cobalt limitation. To demonstrate cobamide sharing by *C. amycolatum*, we employed a co-culture assay using an *E. coli* cobamide auxotroph and show that *C. amycolatum* produces sufficient cobamides to support *E. coli* growth, both in liquid co-culture and when separated spatially on solid medium. We also generated a *C. amycolatum* non-cobamide-producing strain (cob^-^) using UV mutagenesis that contains mutated cobamide biosynthesis genes cobK and cobO and confirm that disruption of cobamide biosynthesis abolishes support of *E. coli* growth through cobamide sharing. Our study provides a unique model to study metabolite sharing by microorganisms, which will be critical for understanding the fundamental interactions that occur within complex microbiomes and for developing approaches to target the human microbiota for health advances.

## Introduction

Microbial communities are complex ecosystems that play essential roles in supporting environmental processes and life on Earth. Fundamental to these microbial systems are the underlying interactions that occur between microorganisms, which can range from cooperative interactions to those that result in competition and predation (1). One broad example of microbial interactions is the process of nutrient sharing and metabolite exchange, which is ubiquitous within microbial communities and can profoundly impact community composition, stability, and function. These metabolites range from metabolic building blocks, such as amino acids, vitamins, and nucleotides, to degradation products of complex polymers. This valuable pool of nutritional resources influences the surrounding community through providing essential nutrients to auxotrophic species and to those that preferentially scavenge the metabolite over costly biosynthesis (2, 3). Discovery and characterization of metabolite exchange is vital for understanding the underlying principles that govern microbial ecology and for manipulating microbial communities for improved medical, agricultural, and environmental outcomes.

Our knowledge of metabolites that influence microbial community dynamics is chiefly based on the “primary economy”, which revolves around carbon flow (4, 5). These reactions are often dependent on essential cofactors, which are small molecules required for catalysis and are considerably less abundant within the ecosystem, but just as important (6). Cobamides, which consist of the vitamin B_12_ family of cofactors, are exceptional compared to other vitamins and cofactors due to their structural diversity and the small number of prokaryotes that can biosynthesize the cofactor. Furthermore, the biosynthetic pathway consists of approximately 30 enzymatic steps, thus making it very energetically costly for the organisms that produce it (7). Many organisms across all domains of life require this cofactor for essential functions that include amino acid metabolism, DNA/RNA synthesis, carbon and nitrogen metabolism, and secondary metabolite synthesis (8, 9). However, *de novo* cobamide biosynthesis is restricted to prokaryotes, indicating that most organisms requiring cobamides must acquire them externally (8). In microbial communities, cobamides are of particular interest because an estimated 86% of bacteria use cobamides, whereas *de novo* cobamide biosynthesis is limited to 37% of bacteria (9). This disproportionate distribution of producers to users, combined with the centrality of cobamides to cellular function and selectivity of microbes towards particular cobamides, suggests that cobamide sharing occurs in microbial communities, and further, that cobamides drive community dynamics (8).

Cobamide sharing has been investigated in diverse ecosystems, including the human gut (6, 10, 11), marine environments (12–14), soil (15), wastewater treatment plants (16), and as we have shown recently, human skin (17). In the gut, approximately half of the known species rely on cobamides for metabolism, and numerous cobamide acquisition mechanisms exist for both cobamide producers and non-producers (18). Furthermore, in various gut models, cobamide supplementation has been shown to lead to differences in microbial community dynamics and to exert protective effects on the host (19–21). Cobamide sharing has been largely studied by genomic analysis, as cobamide biosynthesis, use, and regulation can be effectively predicted from gene content (8).

Previously, we estimated that approximately 39% of species in the skin microbiome encode at least one cobamide-dependent enzyme, whereas only 1% of core skin colonizers are predicted to produce the cofactor *de novo* (17). Furthermore, we found that prevalent skin taxa from the *Cutibacterium* and *Corynebacterium* genus have the potential to produce cobamides, with the relative abundance of cobamide-producing corynebacteria being strongly associated with more diverse skin microbial communities. Across the *Corynebacterium* genus, we also discovered species that produce cobamides *de novo* are almost exclusively host-associated, despite a reduced genome size and the large genetic size of the cobamide biosynthesis pathway. Given the skin microbiome’s essential role in skin health and the limited understanding of the fundamental biological interactions that underlie this contribution, we predict that cobamide sharing occurs on the skin and that this function mediates skin microbiome composition, structure and stability, all of which ultimately influence skin health and barrier function.

To investigate cobamide sharing by skin-associated bacteria, we employed an *in vitro* approach to characterize the ability of diverse *Corynebacterium* species to synthesize cobamides and share them with auxotrophic bacteria. We focus on the validated cobamide producer *C. amycolatum*, showing that this species produces cobamides, regulates biosynthesis as a result of cobalt availability, and shares cobamides with cobamide-dependent species through an undefined mechanism of release. We also generated a cobamide biosynthesis-deficient mutant of *C. amycolatum* and show that the absence of cobamide biosynthesis abates cobamide sharing. Our results highlight the potential for cobamide sharing to influence community dynamics on the skin and underscore the importance of considering ecological interactions within the skin microbiome and their implications for skin health.

## Materials and Methods

### Bacterial strains

*Corynebacterium amycolatum* LK19 (GCA_008042175.1), *Corynebacterium glucuronolyticum* LK488 (GCA_025146395.1), *Corynebacterium kefirresidentii* LK1134 (GCA_025151445.1), and *Corynebacterium jeikeium* LK28 (GCA_014335105.1) are part of the Kalan laboratory skin strain biobank (22). *Corynebacterium glutamicum* ATCC 31269 and *E. coli* ATCC 14169 ( *E. coli* metE^-^) were acquired from the ARS Culture Collection. Skin strain information can be found in Swaney et al. (22). For routine culturing of non-*E. coli* strains, strains were struck out from frozen glycerol stocks onto trypticase soy agar plates supplemented with 0.2% Tween 80 (TSATw80) and incubated at 37°C until colonies of ∼0.5 to 1 mm in size formed. Between 1 and 5 colonies were inoculated into 5 mL brain heart infusion broth with 0.2% Tween 80 (BHITw80) and incubated at 37°C with shaking (200 rpm) for 18-24 h for use in subsequent experiments. *E. coli* strains were cultured on LB plates or in LB broth with overnight incubation at 37°C.

### Illumina+Nanopore hybrid assembly

To generate high-quality complete genome sequences of cobamide-producing species *C. amycolatum* LK19 and *C. glucuronolyticum* LK488, DNA was extracted from 2 mL of overnight cultures using the Sigma Aldrich GenElute Genomic DNA kit following the manufacturer’s instructions, with the following modifications to aid in cell lysis: use of 250 μL of lysozyme solution, addition of 30 μL mutanolysin (4000 U/mL, M9901-5KU) and 37°C incubation at 45 min and 1500 rpm shaking. Long-read sequencing using Oxford Nanopore Technologies (ONT) and hybrid assembly were performed as previously described (23). We did not find any plasmids in the *C. amycolatum* LK19 genome, and the chromosome was circularized and regarded as complete. The *C. glucuronolyticum* LK488 chromosome was circularized and regarded as complete, but there were also five small scaffolds (100 bp to 3.5 kb) that did not circularize.

### Cobamide-related gene and riboswitch analysis

The genomes of *C. amycolatum* LK19, *C. glucuronolyticum* LK488, *C. kefirresidentii* LK1134, *C. jeikeium* LK28, and *C. glutamicum* ATCC 31269 were processed using bakta (v1.8.0) (24) to predict and annotate coding sequences and RNAs. KOfamScan (v1.3.0) (25) was used to scan the genomes for cobamide biosynthesis genes, cobamide-dependent enzymes, and cobamide and cobalt transport proteins (17). The KEGG orthology (KO) identifiers included in the search are indicated in Supplemental Material S1. Hits to the KOs above the predefined inclusion threshold were considered for further analysis. Manual verification using bakta annotations identified the presence of select KOs (*cobC/rnhA, queH, nrdG, prpC, cbiQ, cbiO*) within certain genomes that were not identified via KOfamScan. Sirohydrochlorin ferrochelatase was incorrectly identified in some genomes as *cbiX* (sirohydrochlorin cobaltochelatase); therefore KOfamScan annotation of *cbiX* was also validated manually. As described in Shelton *et al*. (9), presence of the methylcitrate pathway was determined based on the presence of four or more of the following genes: (*prpB, prpC, acnA, acnB, acnD or prpD, sucCD*). Cobalamin riboswitches were identified based on Infernal annotation through the bakta annotation pipeline.

### Cobamide extraction

For cobamide extractions, different synthetic media were used and tailored for optimal growth of each *Corynebacterium* strain. The media used are as follows: CM9 for *C. amycolatum* (17), CM9+1X sweat supplemented with 36 nM Fe(II)SO_4_ for *C. kefirresidentii* and *C. glucuronolyticum*, and CM9+1X sweat supplemented with 36 nM Fe(II)SO_4_ and 1 mM thiamine-HCl for *C. glutamicum* (22). The addition of iron greatly improves growth for the three non-*C. amycolatum* strains when cultured in synthetic media. Additional supplementation with thiamine further promotes growth of *C. glutamicum*, as iron depletion and thiamine synthesis have previously been found to be linked within this species (26).

Strains were cultured overnight in BHITw80. To remove residual cobamides in the medium and scale up culture conditions, cells were washed 3 times with PBS + 0.05% Tween 80, inoculated into 250 mL of the respective synthetic medium for each strain (described above) (starting optical density at 600 nm [OD_600_] = 0.1), and incubated at 37°C shaking (200 rpm) for 24 h. Cells were again washed, inoculated into 1 liter of the respective medium for each strain (OD_600_ = 0.1), and incubated shaking at 37°C shaking for 48 h. Then cobamides were extracted as previously described (17). To measure extracellular cobamide concentration, 1 mL of supernatant was removed from the 1L cultures, sterilized using a 0.2 μm filter, and stored at - 20°C for subsequent analysis.

### E. coli metE^-^ ΔmetH *strain construction*

To generate a control strain for our cobamide indicator assay (17) that would test whether methionine in the sample contributes to growth of the indicator strain *E. coli metE^-^,* we utilized lambda red recombination to construct *E. coli metE^-^* Δ*metH*. This strain only grows with supplemented methionine and therefore indicates the presence of methionine that may confound the estimation of cobamide concentration in the assay.

To prepare electrocompetent *E. coli* ATCC 14169 (metE^-^) for introduction of the red recombinase expression system, 100 mL of LB was inoculated with 2 mL of an overnight, stationary-phase culture of *E. coli metE^-^* and incubated at 37°C until mid-exponential phase was reached (OD_600_=0.3). The culture was then placed on ice for 30 min, followed by centrifugation of 10-mL aliquots for 5 min at 3500 rpm to pellet cells. Cells were washed with 10 mL of ice-cold 10% glycerol four times and resuspended in 100 μL 10% glycerol. In a pre-chilled Eppendorf tube, 1 μL of pKD46 (Supplemental Table 1) and 100 μL of electrocompetent cells were gently mixed and transferred to a pre-chilled electroporation cuvette (0.2 cm gap size). Electroporation was carried out at 2.5 kV (Bio-Rad MicroPulser), and cells were immediately recovered with 1 mL LB and cultured at 28°C for one hour. 100 μL was plated onto carbenicillin plates (100 μg/mL) and incubated at 28°C overnight to select for *E. coli metE^-^* that had taken up pKD46.

The kanamycin resistance cassette was amplified from pKD4 (Supplemental Table 1) using primers with homology extensions that flank the upstream and downstream regions of methionine synthase metH in *E. coli* ATCC 14169 (Supplemental Table 2, homology extensions underlined). The resulting PCR product was purified using the GenElute PCR Clean-Up kit (Sigma-Aldrich #NA1020), digested using DpnI (Promega #R6231), and purified again.

To induce expression of red recombinase, a single colony of *E. coli metE^-^* with pKD46 was inoculated into 2 mL LB with 100 μg/mL carbenicillin and incubated at 28°C until the OD_600_=0.1, at which time 20 μL of 1M L-arabinose was added. Cultures continued to incubate until the OD_600_=0.4-0.6. Preparation of electrocompetent cells were carried out as described above, resuspending the electrocompetent cells in a final volume of 50 μL of 10% glycerol. Electroporation was carried out with 3-5 μL of the linear PCR product (metH homology extension-flanked *kan* cassette) at 2.5 kV (Bio-Rad MicroPulser), and cells were immediately recovered with 1 mL SOC and cultured at 37°C for one hour. 100 μL was plated onto kanamycin plates (20 μg/mL) and incubated at 37°C overnight to select for colonies that had undergone lambda red recombination to replace metH with kan. Successful recombinants were confirmed using PCR primers targeting within (k1, k2, kt) and outside (MetH_Outside_Fwd, MetH_Outside_Rev) (Supplemental Table 2) of the expected locus to test for loss of the parental fragment and gain of the new mutant-specific fragment (27).

The expected phenotype of the new *E. coli metE^-^* Δ*metH* strain was confirmed by culturing both *E. coli metE^-^* and *E. coli metE^-^* Δ*metH* (initial OD_600_=0.01) in 3 mL M9 assay medium (17) for 6 h at 37°C, with or without 0.25 ng/mL cyanocobalamin (CnCbl) or 10 μg/mL methionine. See Supplemental Figure 1 for validation of *E. coli* phenotype.

### Microbiological cobamide indicator assay

The cobamide concentration within cell extracts and supernatants was estimated using the *E. coli* ATCC 14169 (metE^-^) cobamide indicator assay as previously described (17). Minor modifications were made to improve detection, including adjusting the initial *E. coli* OD_600_ to 0.01 and monitoring the OD_600_ of the assay over time to select the optimal time for quantification at which the CnCbl standard curve falls within the linear range. *E. coli metE^-^* Δ*metH* was used as a control strain in the assay to detect methionine and was prepared using the same conditions as *E. coli metE^-^*. CnCbl standards between 0 and 2.5 ng/mL and methionine standards between 0 and 100 μg/mL were added to the assay in a 2.5 μL volume to quantify cobamide concentrations and test for the presence of methionine within the extracts, respectively.

### Cobamide sharing broth co-culture assay

Cells from an overnight BHITw80 culture of *C. amycolatum* LK19 were washed three times with PBS + 0.05% Tween 80, resuspended in CM9 broth, and inoculated into CM9 broth (no CoCl_2_) at an initial OD_600_ of 0.25 for incubation at 37°C shaking (200 rpm) for 24 h. The following day, this *C. amycolatum* pre-culture and *E. coli metE^-^* and *E. coli metE^-^* Δ*metH* overnight cultures were washed three times with PBS + 0.05% Tween 80 and resuspended in CM9 broth. In a 96-well plate (Thermo Scientific #75800-396), 10 µL of each strain was inoculated to a 200 µL total volume of CM9 with or without supplemented 250 nM cobalt, bringing the initial OD_600_ of *C. amycolatum* to 0.15 and *E. coli* strains to 0.015. CnCbl (1 or 2.5 ng/mL) or methionine (10 or 100 µg/mL) were added to cultures in a 2.5 µL volume. To control for the possibility of cobamides in the solution after cell washing, *C. amycolatum* cells from the inoculum were spun down, the supernatant removed and sterilized with a 0.2 μm filter. 10 µL of this solution was added as the filter-sterilized (FS) *C*. amycolatum cell suspension. To test for intracellular cobamides in the inoculum, *C. amycolatum* cells were lysed by performing two cycles of heating the cells at 95°C and freezing at -80°C, followed by 2 minutes of sonication (Branson M2800H ultrasonic bath). 10 µL of this solution was added as the lysed (L) *C. amycolatum* cell suspension. The 96-well plate was then incubated at 37°C stationary for 18 h, after which cultures were serially diluted and plated onto the following medium plates for CFU quantification: LB for *E. coli* mono-cultures, TSATw80 for *C. amycolatum* mono-cultures, LB with 2.5 µg/mL vancomycin to select for *E. coli* from co-cultures, and TSATw80 with 20 µg/mL nalidixic acid to select for *C. amycolatum* from co-cultures. Plates were incubated at 37°C for 24 h for *E. coli* CFU quantification and for 48 h for *C. amycolatum* CFU quantification.

### Cobamide sharing agar co-culture assay

To evaluate cobamide sharing on solid medium, a solid medium with no cobamides or methionine was prepared to support growth of both *C. amycolatum, E. coli* ATCC 14169 (metE^-^), and *E. coli metE^-^* Δ*metH*, termed modified CM9. The composition is identical to CM9 medium (17), but with a reduced MgSO_4_ concentration (0.2 mM) and the addition of 1.5% agarose. The reduced magnesium concentration greatly improves sensitivity of *E. coli* to cobamides, and the use of agarose as opposed to agar prevents undesired *E. coli* growth in the absence of cobamides.

Cells from an overnight BHITw80 culture of *C. amycolatum* LK19 were washed three times with PBS + 0.05% Tween 80, resuspended in 3 mL PBS + 0.05% Tween 80, and adjusted to an OD_600_ of 3.0. A sterile cotton swab (Puritan #25-806-1WC) was used to swab half of a modified CM9 agar plate twice with the cell suspension (round 100 x 15 mm plates with 20 mL volume). In a similar fashion, filter-sterilized and lysed *C. amycolatum* cell suspensions were swabbed onto plates to control for the possibility of cobamides in the solution after cell washing and intracellular cobamides, respectively. CnCbl (10 or 100 ng/mL) or methionine (0.1 or 1 mg/mL) standards were spotted in a 10 µL volume approximately 1 cm from the edge of the center of the plate. Plates were incubated at 37°C for 72 h, allowing for a dense *C. amycolatum* lawn to form. Following, overnight cultures of *E. coli metE^-^* and *E. coli metE^-^* Δ*metH* were washed three times with PBS, resuspended in PBS, and adjusted to an OD_600_ of 0.1. Using a cotton swab, the *E. coli* cell suspensions were struck out in a line from the center of the plate to the outer edge, opposite the side of the *C. amycolatum* lawn, standards, or controls. The plates were incubated for an additional 24 h at 37°C, after which images were taken using a flatbed scanner.

### UV mutagenesis and cobamide biosynthesis screen

From an overnight culture of *C. amycolatum* LK19, 1 mL was pelleted and resuspended in 1mL PBS + 0.05% Tween 80, followed by 30 s of sonication (Branson M2800H ultrasonic bath) to break up cell clumps. 120 μL droplets of cells were placed in a sterile petri dish and exposed to UV light for 7.5 s in a Stratalinker UV UV Crosslinker 2400 (∼4000 µwatts/cm2), previously determined to achieve a death rate between 95-99%. 100 μL cells were then diluted into 5 mL of BHITw80, serially diluted, and plated onto TSATw80 plates to achieve approximately 100-200 colonies per plate. Plates were incubated at 37°C for 48 h to obtain individual colonies.

To screen for *C. amycolatum* mutants that did not produce cobamides, modified CM9 agar plates were prepared (see “Cobamide sharing agar co-culture assay” above for medium composition). The medium was poured in 30-mL volumes into 100 mm x 15 mm square agar plates, which contain thirty-six 13-mm grids (Simport Scientific #D210-16). Single colonies of irradiated *C. amycolatum* were patched onto the modified CM9 grid plates in a 0.5 cm diagonal line. As controls for each plate, one square of the grid was patched with *Corynebacterium jeikeium*, a species that does not produce cobamides, and one was left empty. Cells from an overnight LB culture of *E. coli* ATCC 14169 (*metE^-^*) were washed three times in PBS, adjusted to an OD_600_ to 0.1, and 2 µL spotted within the top left corner of each square. The plates were incubated at 37°C for 18-24 h, after which they were assessed visually for any patches that did not result in *E. coli* growth. Potential mutants were validated by re-streaking the mutant colony onto a fresh modified CM9 plate, incubating at 37°C for 24 h, spotting 10 µL of *E. coli metE^-^* in the adjacent square, and incubating for an additional 24 h.

### Cobamide biosynthesis mutant sequencing variant calling analysis

To validate inactivation of cobamide biosynthesis in *C. amycolatum* cob^-^, genomic DNA was extracted as described in “Illumina+Nanopore hybrid assembly” above. Extracted DNA was sent to SeqCoast (Portsmouth, NH) for Illumina sequencing. Reads were trimmed and processed using fastp (v0.22.0) (28). For alignment to the reference genome, *C. amycolatum* LK19 was first annotated using bakta (v1.8.0) (24), and processed *C. amycolatum* cob^-^ reads were then aligned to the indexed reference genome. Variants were called using freebayes (v0.9.21) (29). Low-quality (q < 30) variants were filtered, and the resulting VCF file was annotated using the bakta *C. amycolatum* LK19 reference genome annotations. Subsequent filtering was performed in R to call high-quality variants (code available on GitHub).

### Protein sequence analysis and structure prediction

For *C. amycolatum* LK19 cob^-^ variants that resulted in missense amino acid substitutions, the protein sequences in which these variants were found were queried against the Uniref50 sequence database using PSI-BLAST using the Phyre2 pipeline (30). Sequence alignments were then visualized with MView (31). Structural prediction of the WT protein was performed using Alphafold2 (32), and Missense3D (33) was used to predict the structural changes introduced by the *C. amycolatum* cob^-^ amino acid substitutions. For cobK and cobO, WT and cob^-^ structures were aligned to the predicted ternary structures of cobK from *Rhodobacter capsulatus* (PDB code: 4X7G) and cobA from *Salmonella enterica* (PDB code: 1G64), respectively, and visualized in PyMOL (v2.5.7).

### HPLC - MS/MS conditions

Cell extracts were analyzed by injecting 5 µL into an Agilent 1290 Infinity II HPLC with an Agilent 6495C iFunnel QQQ mass spectrometer for detection (Agilent, Santa Clara, CA, USA). Separation of the analytes was achieved on an Agilent (Santa Clara, CA, USA) RRHD Eclipse Plus C_18_ (100 mm x 2.1 mm, i.d., 1.8 µm) analytical column using mobile phases consisting of (A) 0.1% (v/v) formic acid in water and (B) 0.1% (v/v) formic acid in acetonitrile and a constant flow of 0.3 mL/min. The autosampler was maintained at 10°C throughout the analysis, and the analytical column was maintained at 30°C. The separation of analytes was achieved by a linear gradient over a run time of 10 min. The HPLC eluent was introduced into the mass spectrometer using electrospray ionization (ESI) in positive mode. The mass spectrometer parameters were drying gas flow rate – 14L/min, drying gas temperature – 250°C, nebulizer pressure – 45 psig, sheath gas flow rate – 12L/min, sheath gas temperature – 300°C, capillary voltage – 3000 V, nozzle voltage – 2000 V, and acquisition rate – 2 cycles/s. Analyte specific transitions used during analysis are as follows: cyanocobalamin [M+2H]^+2^ (678.3 -> 147.1), adenosylcobalamin [M+2H]^+2^ (790.3 -> 147.1), and riboflavin (ISTD) [M+H]^+1^ (383.3 -> 249.0).

### Mass spectrometry quantitative analysis

To evaluate potential matrix effects from the cellular extracts, a post-extraction matrix spike was performed using a set of CnCbl standard dissolved in water at 10, 50, and 500 ng/mL and a set of pooled cellular extract samples from *C. amycolatum* cob^-^ spiked with 10, 50, and 500 ng/mL CnCbl. The samples were analyzed by HPLC-MS/MS using the conditions above, and matrix effects were expressed as a ratio of the mean peak area of the analyte in post-extraction spike samples to the mean peak area of the analyte in standard solutions. We compensated for matrix effects in the analysis by using matrix-matching calibration standards.

The instrument detection limit (IDL) and limit of quantification (LOQ) for CnCbl in water was determined to be 0.32 ng/mL and 1.0 ng/mL, respectively, and the IDL and LOQ for CnCbl in cellular extract was determined to be 1.8 ng/mL and 5.3 ng/mL, respectively. The internal standard used in this analysis was isotope labeled riboflavin-(dioxopyrimidine-^13^C_4_,^15^N_2_). Quantitation of each analyte was based on the peak area measurement. Calibration curves were generated using an unweighted linear regression analysis of the standards and the concentration of each analyte was calculated from the calibration curve. Calibration standards were injected at the beginning and end of each analysis run as well as every 15-20 injections throughout the sequence. The average response of the bracketing calibration standards was used to develop the calibration curve that quantified that set of samples. Suitable sample dilutions were used to ensure the peak areas of samples were within the range of the calibration curve and the dilutions were accounted for in determining the amounts in the samples. All sample injections were made in duplicate and the difference in response was less than ± 10%. The average of the two injections are reported.

## Results

### Cobamide biosynthesis potential and requirements in Corynebacterium species

*De novo* cobamide biosynthesis is encoded by multiple phylogenetic lineages across the *Corynebacterium* genus that include skin-associated species such as *C. amycolatum* and *C. kroppenstedtii* (*17*). We selected four skin isolate *Corynebacterium* strains (*C. amycolatum, C. glucuronolyticum, C. jeikeium,* and *C. kefirresidentii*), as well as the industrially relevant environmental species *C. glutamicum,* for additional investigation into their potential for cobamide biosynthesis and dependence. *C. amycolatum* and *C. glucuronolyticum*, both isolated from umbilicus skin samples, encode *de novo* cobamide biosynthesis genes, with *C. glucuronolyticum* notably missing the *bluB* gene responsible for synthesis of the dimethylbenzimidazole (DMB), the lower ligand of benzimidazolyl cobamides that include cobalamin (Figure 1A). *C. kefirresidentii*, isolated from an occiput skin sample, and *C. jeikeium*, isolated from a skin wound, are both predicted to be non-cobamide-producers, whereas the environmental species *C. glutamicum* is likely a cobamide salvager because it encodes *pduO*, *cobP/cobU*, *cobS/cobV, cobU/cobT, and cobC*, which are involved in cobinamide salvaging (34).

**Figure 1.**
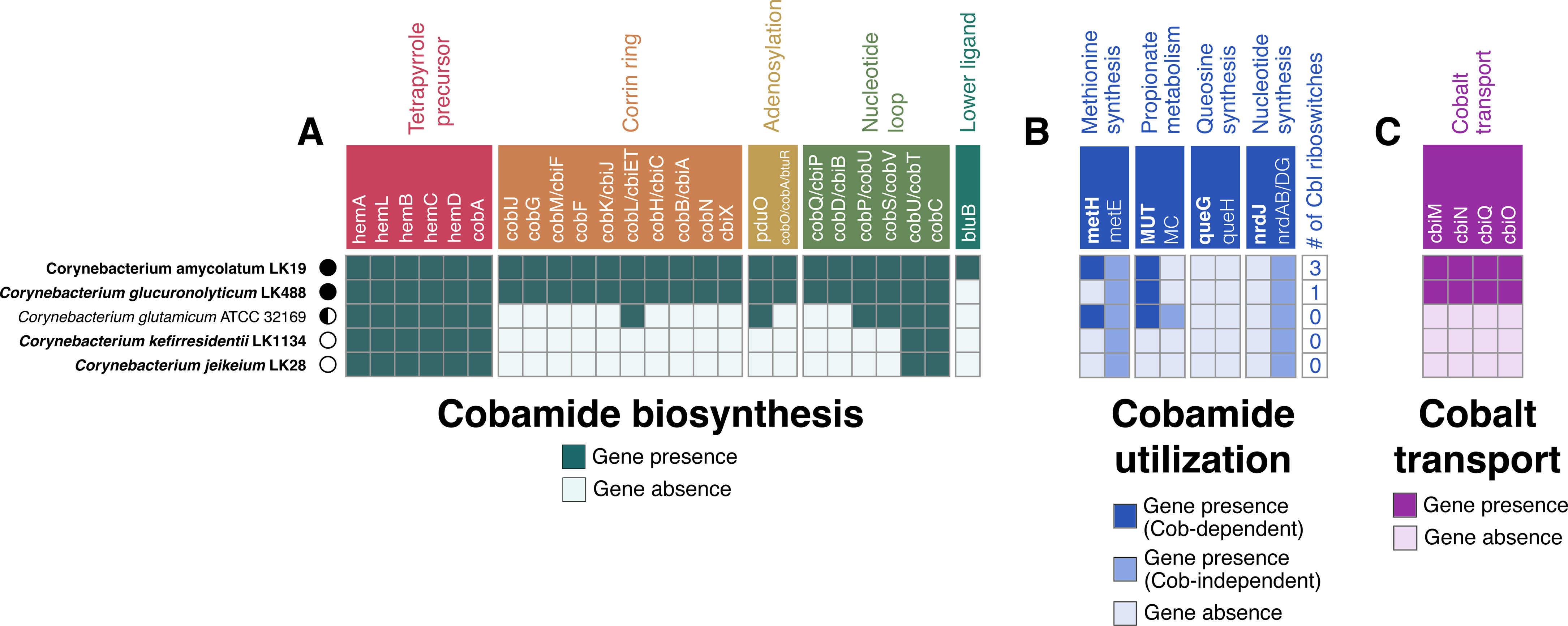
Cobamide biosynthesis and dependence of bacterial strains used in this study. The genomes of strains used in this study were scanned for (A) cobamide biosynthesis genes, (B) cobamide-dependent and -independent genes, and (C) cobalt transport genes using KOFamScan. Cobalamin riboswitches were identified within the genomes using Infernal through the Bakta annotation pipeline. Strains isolated from human skin are shown in bold italics. Cobamide biosynthesis genes in A) are differentially colored based upon their cobamide biosynthesis subsection. Genes encoding for cobamide-dependent enzymes in B) are bolded in white, and dark blue squares indicate gene presence, whereas the intermediate-colored blue squares indicate gene presence of their cobamide-independent isoforms. MC = methylcitrate pathway. Cobamide biosynthesis prediction for each strain is represented as follows: producer = black circle, salvager = semi-circle, non-producer = white circle.

Prokaryotes carry out cobamide biosynthesis as a means to facilitate various enzymatic reactions that are reliant on cobamides for essential cofactor activity (9). We thus examined the five *Corynebacterium* genomes for cobamide-dependent enzymes, their cobamide-independent isoforms, and cobalamin riboswitches, which are regulatory RNA elements that regulate nearby genes related to cobamide metabolism in response to cobamide concentration (35–37). *C. kefirresidentii* and *C. jeikeium* lack cobamide-dependent enzymes and cobalamin riboswitches (Figure 1B), consistent with their inability to biosynthesize cobamides. In contrast, *de novo* producers *C. amycolatum* and *C. glucuronolyticum* exhibit distinct cobamide metabolic requirements. *C. amycolatum* encodes two cobamide-dependent enzymes (methionine synthase and methylmalonyl coA-mutase) and three cobalamin riboswitches regulating cobalt transport and cobamide-independent methionine synthesis. *C. glucuronolyticum* encodes only methylmalonyl coA-mutase and a single riboswitch upstream of a cobalt transport locus. The salvager *C. glutamicum* has enzymes for both cobamide-dependent and -independent methionine synthesis and propionate metabolism, but lacks cobalamin riboswitches.

Central to cobamide biosynthesis is the insertion of the trace element cobalt into the corrin ring, thus necessitating the scavenging of cobalt from the environment using high-affinity cobalt uptake systems (38). When scanned for the *cbiMNQO* cobalt transport system, both *C. amycolatum* and *C. glucuronolyticum* were found to encode these genes, whereas the remaining *Corynebacterium* species did not (Figure 1C). Overall, we find that the potential for cobamide biosynthesis across phylogenetically distinct *Corynebacterium* species is reflected through their metabolic dependence upon cobamides (i.e., the presence of cobamide-dependent enzymes and cobalamin riboswitch regulation) and their ability to transport cobalt.

To validate the genomic prediction of cobamide biosynthesis potential of the *Corynebacterium* species, we cultured the two predicted producers *C. amycolatum* and *C. glucuronolyticum,* the predicted salvager *C. glutamicum,* and the predicted non-producer *C. kefirresidentii* in cobamide-free minimal medium and then extracted cobamides from the cells for quantification in an *E. coli* microbiological indicator assay (17). Supporting genomic prediction, *C. amycolatum* and *C. glucuronolyticum* both produce cobamides, with *C. amycolatum* producing an estimated 30-fold higher amount per gram of wet cell weight when compared to *C. glucuronolyticum* under similar growth conditions (1.51 μg vs. 0.05 μg respectively, with 50 nM supplemented CoCl_2_) (Figure 2). Because coordination of cobalt within the corrin ring of cobamides is essential for biosynthesis (38), we tested cobamide production by *C. amycolatum* under low (0 nM) and high (250 nM) cobalt concentrations. This revealed that cobamide biosynthesis within this species is highly dependent upon and regulated by cobalt concentration (0.08 vs. 4.58 μg cobamides per gram wet cell weight respectively). Mass spectrometry validated these findings, demonstrating an average of 0.017 and 1.17 μg cobamide produced per gram wet cell weight in 0 nM vs 250 nM CoCl_2_, respectively, and identified that the specific cobamide produced by *C. amycolatum* is cobalamin (Supplemental Table 3, Supplemental Figure 2). Interestingly, the concentration of cobalamin in *C. glucuronolyticum* cell extracts was below the limit of quantification (LOQ, 5.3 ng/mL) by mass spectrometry, even though the predicted cobamide concentration using the indicator assay was well above this value (54.7 ng/mL). Because the indicator assay is responsive to cobamides with different lower ligand structures, this suggests that *C. glucuronolyticum* may produce cobamides with a non-benzimidazolyl lower ligand, supporting the finding that this species does not encode the *bluB* gene responsible for DMB synthesis.

**Figure 2.**
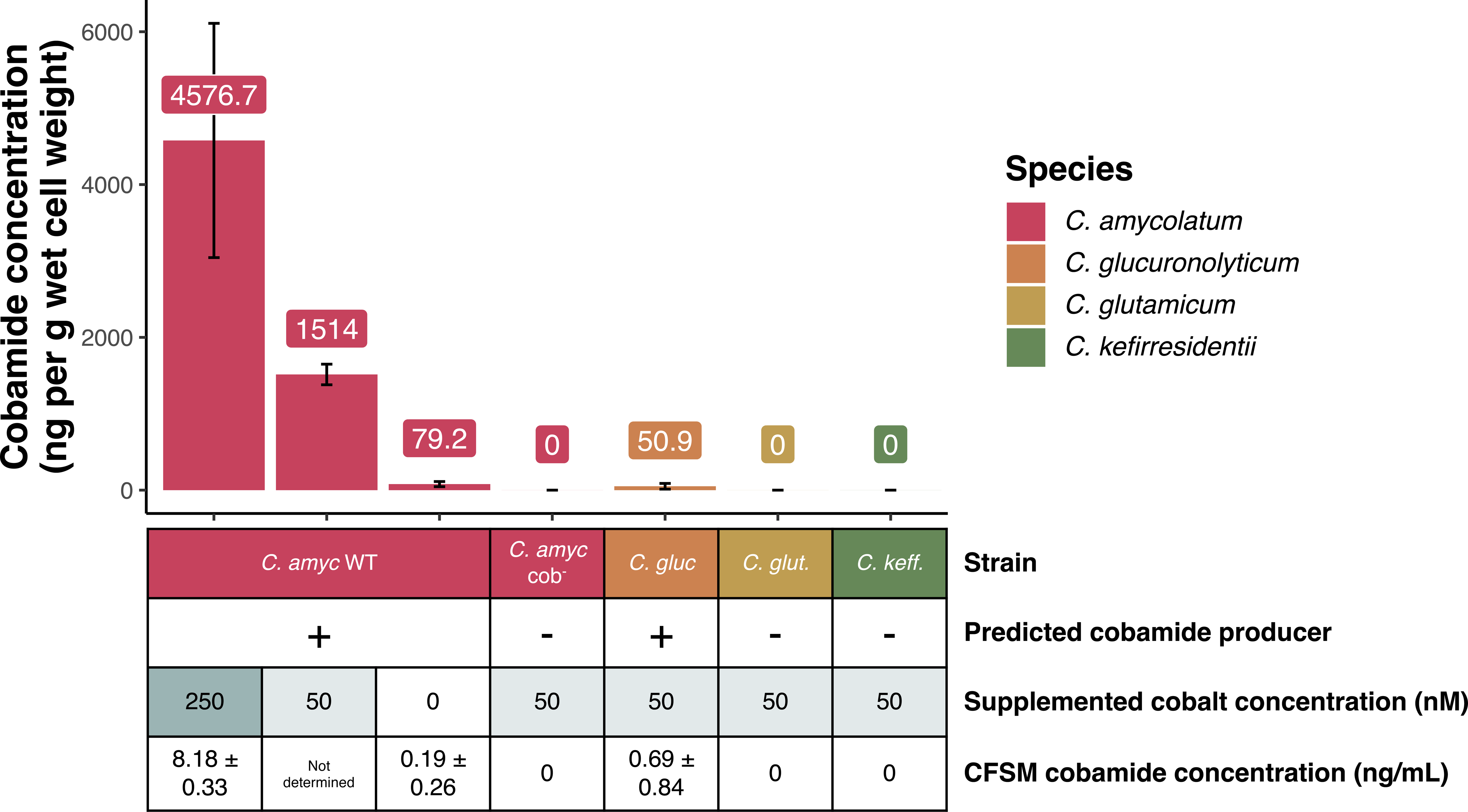
Cobamide biosynthesis is carried out by *Corynebacterium* predicted producers and is restricted by cobalt availability. *C. amycolatum* WT, *C. amycolatum* cob-, *C. glucuronolyticum, C. glutamicum,* and *C. kefirresidentii* were cultured in minimal cobamide-free medium with 0, 50, or 250 nM CoCl_2_ concentrations. Intracellular and extracellular cobamides were estimated using an *E. coli* microbiological indicator assay (17). Cobamide vs. methionine presence in the samples was confirmed using both the *E. coli metE-* and *E. coli metE^-^* Δ*metH* indicator strains. Intracellular cobamides were quantified per gram of wet cell weight, and extracellular (CFSM) cobamides were quantified per mL of media. Intracellular cobamides were converted to their cyano form before testing in the indicator assay, and extracellular cobamides were tested in their native form.

Cobamides were not detected in *C. glutamicum* or *C. kefirresidentii* cellular extracts by the indicator assay method, validating the genomic prediction that they are not capable of synthesizing cobamides. Cobamides were also present within the cell-free spent medium of *C. amycolatum* (8.18 ng/mL in 250 nM CoCl_2_) and *C. glucuronolyticum* (0.7 ng/mL in 50 nM CoCl_2_), determined using the indicator assay, supporting the possibility of cobamide sharing by these two skin species (Figure 2).

### C. amycolatum shares cobamides with E. coli cobamide auxotroph

To evaluate the ability of *C. amycolatum* to share cobamides, we developed a co-culture system using *E. coli metE^-^,* whose growth requires cobamides or methionine (Figure 3A), and control strain *E. coli metE^-^* Δ*metH*, whose growth requires methionine (Figure 3B), to assess whether *C. amycolatum* could support growth of the cobamide auxotroph *E. coli metE^-^* in co-culture. We tested this system under two defined cobalt conditions, low cobalt (0 nM CoCl_2_) and high cobalt (250 nM CoCl_2_), as this should limit cobamide biosynthesis by *C. amycolatum* because of the essentiality of cobalt for cobamide production (38). In this co-culture system, *C. amycolatum* WT produces sufficient cobamides to support *E. coli metE^-^*growth, with a 1.4-log_10_ increase in *E. coli metE^-^* CFUs under high cobalt conditions when compared to low cobalt conditions (Figure 3C). Minimal growth of the control strain *E. coli metE^-^* ΔmetH in *C. amycolatum* WT co-culture supports that cobamides, rather than methionine, are responsible for the observed *E. coli metE^-^* growth response (Figure 3D). The ∼0.5 log_10_ increase in growth of *E. coli metE^-^* when co-cultured with *C. amycolatum* WT under low cobalt suggests that there is likely trace amounts of residual cobalt available to *C. amycolatum* for cobamide biosynthesis.

**Figure 3.**
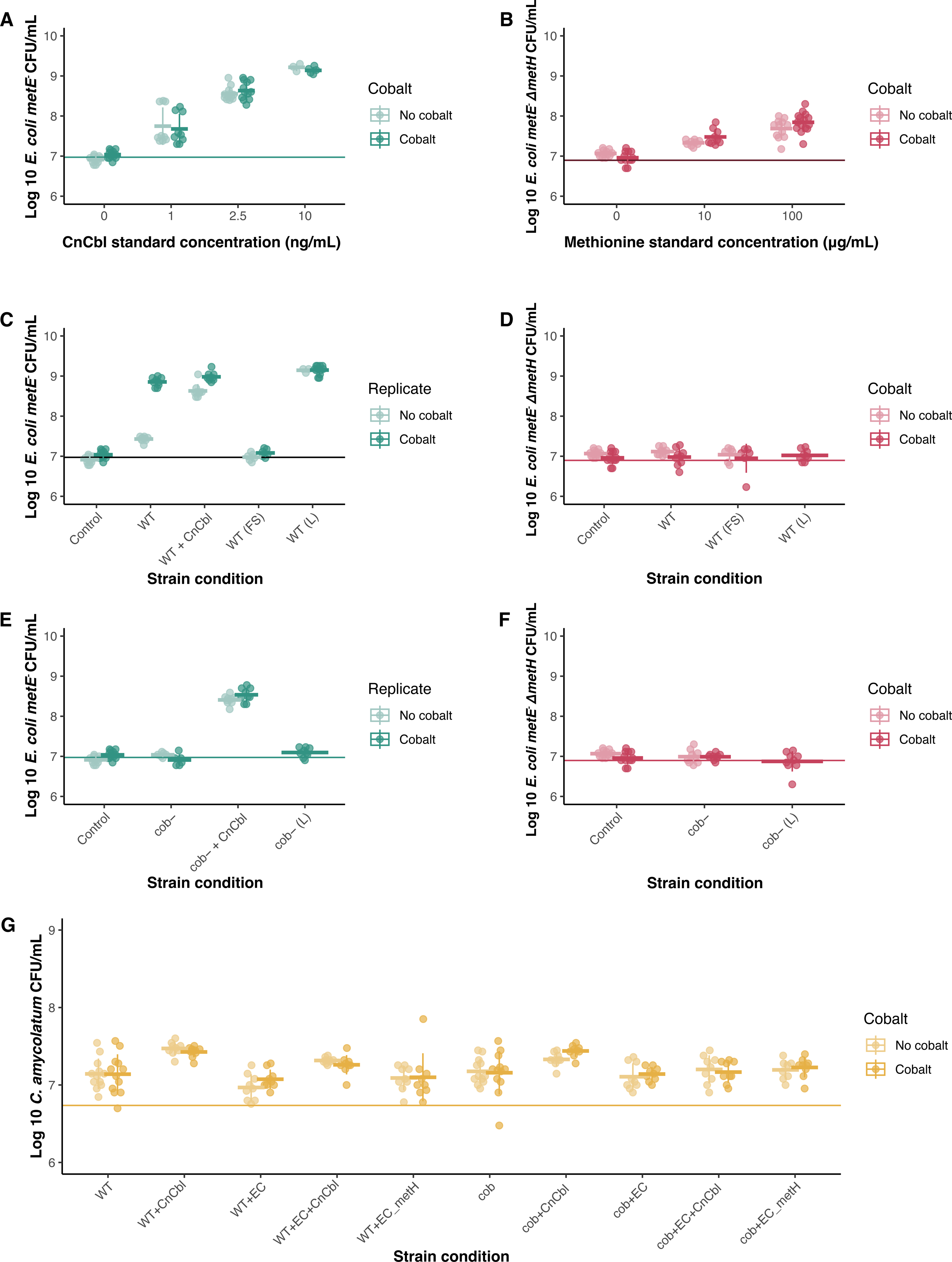
*C. amycolatum* shares cobamides with cobamide auxotroph and regulates cobamide biosynthesis through cobalt availability. **(A)** *E. coli metE^-^* (blue) was cultured with 0, 1, and 2.5 ng/mL cyanocobalamin standards (CnCbl), and **(B)** *E. coli metE^-^* Δ*metH* (pink) was cultured with 0, 10, and 100 μg/mL methionine to validate expected growth response of each strain, quantified in CFU/mL. Growth of *E. coli metE^-^* is proportional to the concentration of cobamides or methionine in the medium, and growth of *E. coli metE^-^* Δ*metH* is proportional to the concentration of only methionine, thus distinguishing between cobamides and methionine in supporting *E. coli* growth. *C. amycolatum* WT (WT) was co-cultured in minimal medium with **(C)** *E. coli metE^-^*(blue) or **(D)** *E. coli metE^-^* Δ*metH* (pink) under low (0 nM) or high (250 nM) cobalt concentrations. *C. amycolatum* cob^-^ (cob-) was co-cultured in minimal medium with **(E)** *E. coli metE^-^*(blue) or **(F)** *E. coli metE^-^* Δ*metH* (pink) under low (0 nM) or high (250 nM) cobalt concentrations. **(G)** *C. amycolatum* CFU/mL was measured to demonstrate growth across each strain condition. n=9 replicates for each condition (3 technical replicates across 3 biological replicates). Solid lines represent the measured inoculum for *E. coli metE^-^*, *E. coli metE^-^* Δ*metH*, or *C. amycolatum* averaged across all replicates. FS = filter-sterilized *C. amycolatum* cell suspension, L = lysed *C. amycolatum* cell suspension.

Furthermore, supplementation of *C. amycolatum* WT co-culture with 2.5 ng/mL CnCbl rescues *E. coli metE^-^* growth (∼1.2 log_10_ increase) under low cobalt conditions and that *C. amycolatum* WT continues to synthesize and share cobamides even when provided with the vitamin exogenously, producing comparable *E. coli metE^-^* growth under high cobalt conditions vs low cobalt conditions with supplemented CnCbl (8.9 vs 9.0 log_10_ CFU/mL, respectively). Lastly, filter-sterilized *C. amycolatum* WT cell suspension failed to yield growth of *E. coli metE^-^*, whereas lysed *C. amycolatum* WT cells supported robust growth of the indicator strain, supporting that the cobamides being shared during co-culture are from intracellular cell contents rather than extracellular residual cobamides from the medium (Figure 3C). Additionally, we show that *C. amycolatum* exhibits similar growth across all conditions, demonstrating that differences in *E. coli* growth are a result of differential *C. amycolatum* cobamide biosynthesis rather than differential *C. amycolatum* growth (Figure 3G). Altogether, our findings demonstrate the ability of *C. amycolatum* to share cobamides with other bacterial species, that *C. amycolatum* cobamide biosynthesis is tightly regulated as a result of cobalt availability, and that cobamide biosynthesis and sharing is carried out even with exogenous cobamide supplementation.

In the liquid medium co-culture system, *C. amycolatum* and *E. coli* occupy the same physical space and come into very close cell-cell contact, facilitating nutrient exchange. To determine whether *C. amycolatum* could produce and share cobamides when separated physically from *E. coli*, we cultured *C. amycolatum* on minimal medium plates for 3 days to allow for dense growth and biosynthesis of cobamides, after which *E. coli metE^-^* or *E. coli metE^-^* Δ*metH* was inoculated onto the opposite side of the plate. Results show that *C. amycolatum* WT does indeed support growth of *E. coli metE^-^*, but not *E. coli metE^-^* Δ*metH,* confirming the ability of *C. amycolatum* to produce and share cobamides (Figure 4A-C). Similar to liquid culture, we show that cobalt availability regulates cobamide biosynthesis and sharing by *C. amycolatum* WT on solid medium, where a dramatic reduction in *E. coli metE^-^* growth is observed under low cobalt (0 nM cobalt) when compared to high cobalt (250 nM cobalt) conditions (Supplemental Figure 3). Furthermore, filter-sterilized *C. amycolatum* WT cell suspension did not yield growth of *E. coli metE^-^*, whereas lysed *C. amycolatum* WT cells supported slight growth (Figure 4A, G-H), demonstrating that intracellular *C. amycolatum* cobamides are shared with *E. coli metE^-^*, as opposed to residual cobamides or methionine present in the cell solution. Validating the expected growth response of *E. coli metE^-^*and *E. coli metE^-^* Δ*metH* to cobalamin and methionine standards, we show that neither strain grows in the absence of cyanocobalamin or methionine, *E. coli metE^-^*responds to both cyanocobalamin and methionine, and *E. coli metE^-^*Δ*metH* responds only to methionine (Figure 4A, I-L). In all, these findings further support our hypothesis that *C. amycolatum* produces and shares cobamides with other bacteria. Furthermore, the presence of *E. coli* growth in a spatially distinct location from *C. amycolatum* in solid-phase culture suggests that cobamides are secreted or released by *C. amycolatum*, although additional experiments are necessary to determine the mechanism of release.

**Figure 4.**
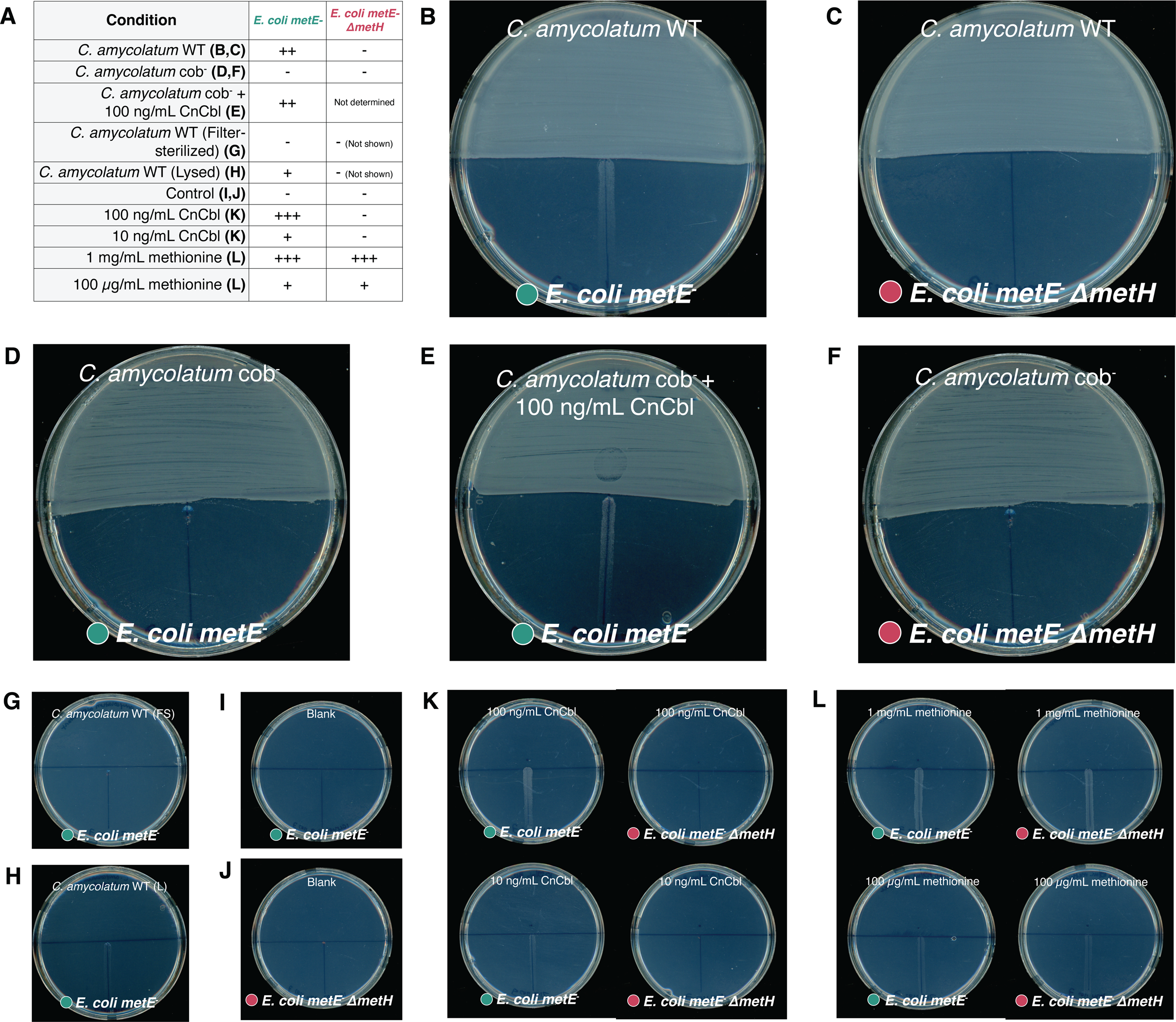
*C. amycolatum* releases cobamides on solid medium to support growth of spatially separated cobamide auxotroph *E. coli metE^-^.* *C. amycolatum* WT or cob^-^ cell suspension or controls (CnCbl, methionine, filtered cell suspension (FS), lysed cell suspension (L) were added to half of a minimal medium plate and incubated for 3 days, after which *E. coli metE^-^*(blue) or *E. coli metE^-^* Δ*metH* (pink) was struck out on the adjacent side of the plate. *E. coli* growth was recorded after 24 h. Growth summary for all conditions is indicated in **(A).** - indicates no growth and +, ++, +++ indicate increasing levels of growth, respectively. *C. amycolatum* WT lawn with **(B)** *E. coli metE^-^* and **(C)** *E. coli metE^-^* Δ*metH.* C. amycolatum cob^-^ lawn with **(D)** *E. coli metE^-^,* **(E)***E. coli metE^-^* + 100 ng/mL CnCbl, and **(F)** *E. coli metE^-^* Δ*metH. E. coli metE^-^* with **(G)** *C. amycolatum* filter-sterilized cell suspension and **(H)** *C. amycolatum* lysed cell suspension. Blank controls for **(I)** *E. coli metE^-^* and **(J)** *E. coli metE^-^* Δ*metH*. **(K)** *E. coli metE^-^* and **(L)** *E. coli metE^-^* Δ*metH* growth response to CnCbl and methionine standards.

### C. amycolatum cob^-^ does not produce detectable cobamides

To produce a cobamide biosynthesis deficient strain of *C. amycolatum*, we attempted to delete the cobGHIJ operon using a suicide plasmid-mediated homologous recombination approach.

However, we found that this species is particularly resistant to the introduction of foreign DNA. Therefore, we employed UV mutagenesis to induce random mutations within the *C. amycolatum* genome in combination with a screen based on the solid-phase cobamide sharing assay to identify non-cobamide-producing *C. amycolatum* mutants. The screen utilizes *E. coli metE^-^* as an indicator of cobamide presence in the medium. Therefore, we tested *C. amycolatum* colonies mutagenized with UV light on minimal medium for their inability to support *E. coli metE^-^* growth (Supplemental Figure 4A). With this approach, we screened ∼1800 colonies and isolated a single mutant of *C. amycolatum*, which we designate *C. amycolatum* cob^-^, that does not produce detectable cobamides when cultured in minimal medium and assayed with the *E. coli metE^-^* indicator assay and mass spectrometry (Figure 2, Supplemental Figure 4B, Supplemental Table 3).

Whole genome sequencing and variant calling of *C. amycolatum* cob^-^ revealed two distinct SNPs in two cobamide biosynthesis genes; cobK (precorrin-6A reductase, involved in corrin ring synthesis) and cobO (corrinoid adenosyltransferase, involved in adenosylation) (Supplemental Figure 4C). There were 16 total predicted variants, 10 of which occurred within coding regions, and of these 7 that resulted in amino acid changes, including cobK and cobO (Supplemental Material S2). The 5 other off-target SNPs were located in the following genes: murA (UDP-N-acetylmuramate dehydrogenase, involved in peptidoglycan biosynthesis), miaB (tRNA (N6-isopentenyl adenosine(37)-C2)-methylthiotransferase, involved in tRNA modification of isopentenylated adenosine-37), pepN (aminopeptidase N), DUF3329 domain-containing function (unknown function), and a 4Fe-4S dicluster domain protein (unknown function).

A PSI-BLAST alignment of diverse homologs for both cobK and cobO revealed that the amino acid changes occurred in highly conserved regions of both proteins (Figure 5A,C). Furthermore, structural prediction of the mutant protein sequence using AlphaFold2 and Missense3D suggests that the cobK mutation results in an altered cavity. To assess how the mutation may impact substrate or cofactor binding, we aligned the predicted structure of *C. amycolatum* cobK to the ternary structure of cobK from *Rhodobacter capsulatus* (*39*), which revealed that the change from arginine to cysteine potentially disrupts the precorrin-binding region (Figure 5B). Similarly, alignment of *C. amycolatum* cobO to the predicted ternary structure of cobA from *Salmonella enterica* (*40*) suggests that the S44F mutation likely alters the P-loop and ATP-binding region, as the mutation occurs within the serine residue of the P-loop motif (Figure 5D). This is a glycine-rich sequence followed by a conserved lysine and a serine or threonine that is important for ATP binding (41). Missense3D did not predict any structural changes to cobO as a result of the mutation, however. The remaining five off-target mutations did not occur in conserved regions nor did they have predicted structural changes (Supplemental Figure 5, Supplemental Material S2).

**Figure 5.**
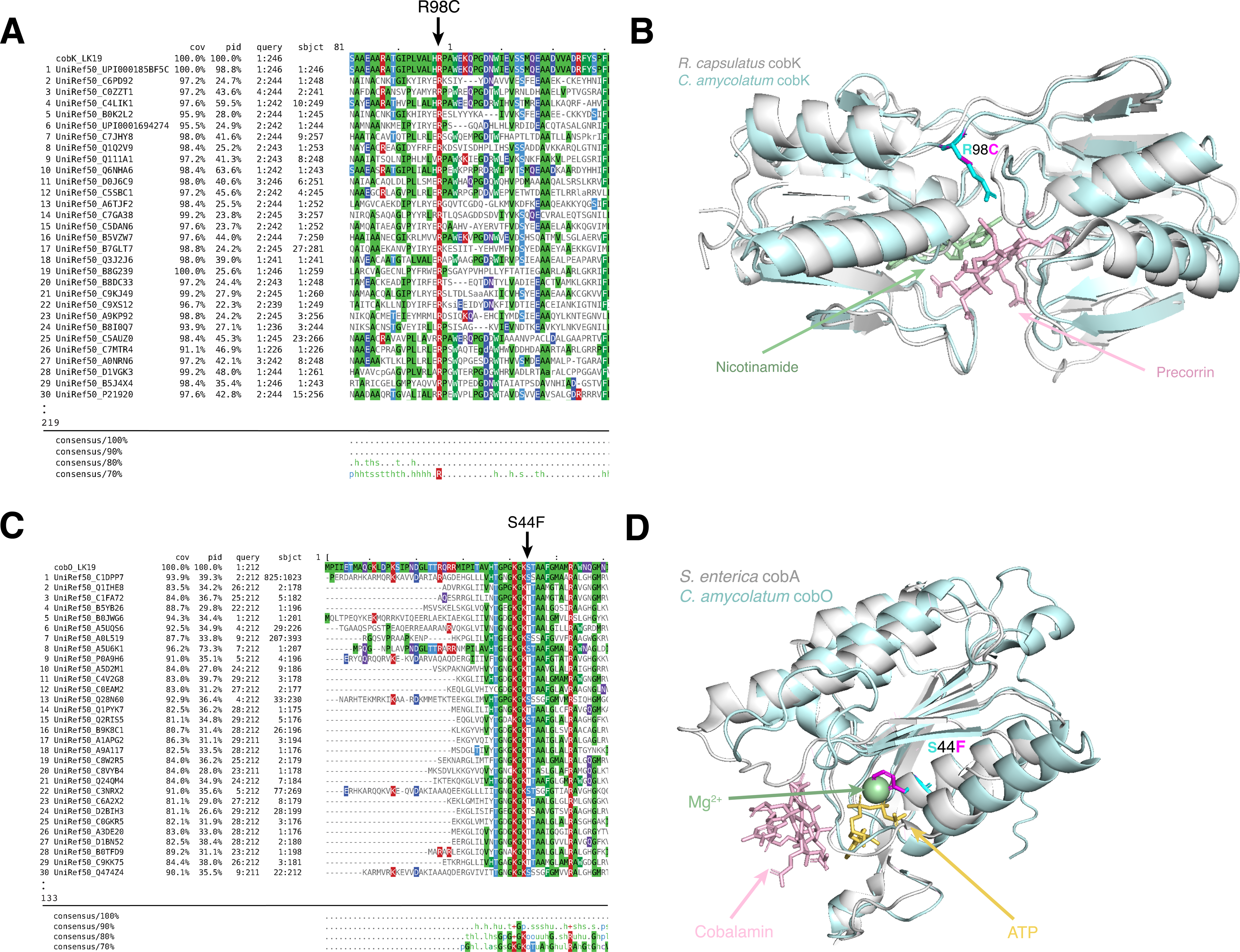
Mutation of cobK and cobO in a non-cobamide producing strain of *C. amycolatum.* Sequence homologs of **(A)** cobK and **(C)** cobO were detected by querying *C. amycolatum* WT protein sequences against the Uniref50 sequence database using PSI-BLAST through the Phyre2 pipeline. Sequences are visualized with MView, with residues colored by identity. The top 30 alignments are shown. The consensus sequence is based on all alignments (up to 1000). Alphafold2 was used to predict the structure of **(B)** cobK and **(D)** cobO. Structures were then aligned to the predicted ternary structures of cobK from *Rhodobacter capsulatus* (PDB code: 4X7G) and cobA from *Salmonella enterica* (PDB code: 1G64). *C. amycolatum* WT structures are shown in light blue and PDB structures are shown in white. Missense3D was then used to predict the structure of *C. amycolatum* cob^-^ cobK and cobO. Residue changes are shown in **(B)** for cobK and **(D)** for cobO, with the wild-type residue depicted in cyan and the cob^-^ residue in magenta.

We next compared morphological and growth characteristics of *C. amycolatum* WT to cob^-^ in rich media and minimal media. Morphologically, *C. amycolatum* cob^-^ colonies were indistinguishable from WT, both similar in size and shape, when grown on rich medium (Supplemental Figure 6A-B). Similarly, both strains exhibited similar growth in rich and minimal liquid broth, characterized by autoaggregation, ring formation on the culture tube, and settling of cells at the tube bottom (Supplemental Figure 6C-D). A slight increase of suspended, non-aggregated cells was observed for *C. amycolatum* cob^-^ in certain medium types (Supplemental Figure 6C). The two strains exhibited very similar growth curves in a range of medium types at low and high starting cell densities (Supplemental Figure 6E); however in rich BHI + 0.2% Tween 80 medium, a slight increase in lag time can be observed for *C. amycolatum* cob^-^. Overall, we find that *C. amycolatum* cob^-^, while being deficient in cobamide biosynthesis, does not demonstrate strong growth defects or differences when compared to *C. amycolatum* WT in the media tested.

Lastly, we tested cobamide sharing by *C. amycolatum* cob^-^, finding that this cobamide-deficient strain failed to support growth of *E. coli metE^-^* in co-culture or after lysing the cell suspension inoculum (Figure 3E-F). Similarly, no cobamide sharing was observed on solid medium by *C. amycolatum* cob^-^ (Figure 4D-F). In both assays, addition of CnCbl to the system restored growth of cobamide auxotroph *E. coli metE^-^* (Figure 3E and Figure 4E). Overall, these findings reinforce that *C. amycolatum* cob^-^ is unable to synthesize cobamides and lends additional evidence to the cobamide sharing nature of *C. amycolatum*.

## Discussion

Our study provides the first report of cobamide sharing by skin-associated bacteria and demonstrates the variable cobamide biosynthetic potential across different species of the *Corynebacterium* genus. Through *in vitro* culturing of phylogenetically distinct *Corynebacterium* species, we show that species predicted to encode cobamide biosynthesis do produce cobamides and even share them with other cobamide-dependent bacteria through an unknown mechanism of release. Furthermore, we show that *C. amycolatum* deficient in cobamide biosynthesis harbors mutations within cobK and cobO, which either together or individually result in disruption of *de novo* cobamide biosynthesis in *C. amycolatum,* providing a valuable tool for probing cobamide-mediated interactions in skin-associated bacteria.

In this study, we illustrate the power of genomics to predict function in the *Corynebacterium* genus. We show that we can accurately predict cobamide biosynthesis potential in diverse *Corynebacterium* species through assessing cobamide biosynthesis pathway presence, cobamide-dependent metabolism, and cobalt transport potential. In *C. amycolatum* and *C. glucuronolyticum*, both species encode *de novo* cobamide biosynthesis, cobalt transport, and cobamide-dependent metabolism, yet exhibit vastly different cobamide production levels. Thus, this finding highlights a key limitation of using genomics to predict functional outcomes, which is that we lack information about the downstream processes of gene expression, protein expression, and protein activity that result in the metabolic end product. Furthermore, a notable difference between the two species is that in addition to encoding for methylmalonyl coA-mutase, *C. amycolatum* also encodes cobamide-dependent methionine synthase. This could suggest that the increased production of cobamides may be a result of an increased metabolic need for the cofactor and may also suggest differences in cobamide-related gene or protein regulation. Alternatively, the differences may lie in the exact cobamide synthesized by *C. glucuronolyticum*, which does not encode *bluB* (5,6-dimethylbenzimidazole synthase), responsible for synthesis of benzimidazolyl cobamides. This suggests it may produce a cobamide with another lower ligand, such as purinyl or phenolyl cobamides. Indeed, *C. glucuronolyticum* cell extract supported *E. coli metE^-^* growth in the indicator assay, demonstrating the presence of cobamides. Yet, mass spectrometry revealed very low levels of cobalamin (a benzimidazolyl cobamide) in the same cell extracts, supporting this prediction.

A wealth of studies exist that predict the existence of cobamide sharing between microorganisms (9, 11–17, 42), and of these, a number provide direct experimental evidence for this phenomenon, although they are almost exclusively restricted to those between bacteria and cobamide-requiring eukaryotes (43–46). Evidence of cobamide sharing between two or more bacterial species is limited, potentially due to a lack of experimental models to measure this phenomenon and the non-auxotrophic nature of many cobamide-utilizing bacteria (9). Existing examples highlight the importance of cobamide precursor salvaging and remodeling, such as within dechlorinating systems (47–49), among amoeba-bacterial models (50, 51), and in synthetic cobamide salvaging models (52). Our study introduces a co-culture system featuring *C. amycolatum,* a cobamide-producing skin-associated bacteria, and *E. coli metE^-^*, a cobamide auxotroph. Our model has demonstrated cobamide release by *C. amycolatum* supporting *E. coli metE^-^*growth, both in a mixed broth co-culture and on spatially separated solid medium. The mechanism behind cobamide release from *C. amycolatum* remains unclear, as no active cobamide export mechanisms have been identified in bacteria (8). Possibilities include cell death, a ubiquitous process that is predicted to supply nutrients to neighboring live cells (53), or cell lysis, which can occur naturally through mechanisms such as exposure to toxins or through bacteriophage infection. Indeed, lysis mediated by bacteriophages has been demonstrated to release intracellular amino acids to support robust growth of amino acid auxotrophic bacteria, more so than mechanical lysis or secretion (54). Therefore, similar mechanisms may exist for cobamide release and provisioning to the community by *C. amycolatum*.

We generated a cobamide biosynthesis-deficient strain of *C. amycolatum*, which we show has mutations within cobK and cobO. Mutations and deletions of cobK and cobO in other bacteria or archaea have been previously identified to disrupt cobamide biosynthesis or growth under cobamide-dependent conditions (55–60), demonstrating that these genes are critical for cobamide biosynthesis. Curiously, we were only able to isolate one cobamide biosynthesis-deficient mutant strain of *C. amycolatum*, even though the cobamide biosynthesis pathway makes up approximately 1% of the genome, and the isolation of two independent mutations should be rarer than one single mutation. It is unknown whether the occurrence of mutations within two separate biosynthesis genes is merely a coincidence or if rather this was selected for as a consequence of the design of the mutagenesis screen. Considering that the mutations both occur within highly conserved regions of the protein sequences and are predicted to occur within close proximity to critical binding sites, it is plausible that both mutations do disrupt protein function and that both disruptions were necessary for identification of this specific *C. amycolatum* mutant, although the exact mechanism for this is unknown.

*C. amycolatum* cob^-^ displays minor growth and autoaggregation deficiencies compared to WT when cultured in certain medium types, likely linked to altered methionine and propionate metabolism resulting from the inactivated cobamide biosynthetic pathway. Indeed, cobamide-independent methionine synthase metE in other organisms is predicted to be less efficient (61), thermally sensitive (62, 63), and resource-inefficient (64) when compared to cobamide-dependent metH. However, because *C. amycolatum* does not encode the cobamide-independent pathway for propionate metabolism as it does for methionine synthase (Figure 1B), we thus postulate that the inability to produce cobamides may significantly alter *C. amycolatum* propionate metabolism, as accumulation of propionate and propionate-derived catabolites can be toxic to bacteria (65). Interestingly, methylmalonyl-CoA, an intermediate of propionate metabolism, is incorporated into cell wall lipids in *M. tuberculosis* as a means to detoxify propionate metabolites (66). Therefore, we predict that in *C. amycolatum* cob^-^, propionate metabolism is disrupted as a result of the inability to use the cobamide-dependent methylmalonyl pathway, leading to the accumulation of toxic metabolites and the shunting of methylmalonyl-CoA towards cell wall lipid synthesis in an effort to detoxify these metabolites. This provides a possible explanation for why *C. amycolatum* cob^-^ exhibits slight differences in growth and autoaggregation compared to WT. Alternatively, we cannot rule out the possibility that the off-target mutations from UV mutagenesis contribute to the observed defects.

In conclusion, we provide an *in vitro* model to interrogate cobamide sharing by skin commensals and provide evidence that a skin-associated member of the *Corynebacterium* genus can share cobamides with other bacteria. Our study advances the understanding of the role of corynebacteria on the skin and adds to a growing list of functions that members of this previously understudied taxon perform in the skin environment (67–70). Furthermore, generation of a cobamide biosynthesis-deficient mutant of *C. amycolatum* reveals key mutations that likely abrogate biosynthesis and provides a useful model for studying the effects of cobamide biosynthesis and sharing among skin-associated bacteria. Future studies to identify other shared nutrients within the skin environment and to apply this system to skin microbial community models will provide important insight into how microbial metabolites drive community dynamics and ultimately modulate host health.

## Data Availability

Genomes for skin-associated bacterial strains used in this study are deposited under BioProjects PRJNA395377 and PRJNA803478.

## Funding and acknowledgements

This work was supported by grants from the National Institutes of Health (NIAID U19AI142720 [L.R.K], NIGMS R35 GM137828 [L.R.K], and NIAMS F31AR079846 [M.H.S.]. The content is solely the responsibility of the authors and does not necessarily represent the official views of the National Institutes of Health. The authors gratefully acknowledge the following individuals: Rauf Salamzade for assistance with hybrid genome assembly, Michael Thomas for providing E. coli cloning strains, and members of the Kalan Laboratory for feedback and discussion.

## References

1. Faust K, Raes J. 2012. Microbial interactions: from networks to models. Nat Rev Microbiol 10:538–550.

2. Giri S, Oña L, Waschina S, Shitut S, Yousif G, Kaleta C, Kost C. 2021. Metabolic dissimilarity determines the establishment of cross-feeding interactions in bacteria. Curr Biol 10.1016/j.cub.2021.10.019.

3. Pierce EC, Dutton RJ. 2021. Putting microbial interactions back into community contexts. Curr Opin Microbiol 65:56–63.

4. Rowland I, Gibson G, Heinken A, Scott K, Swann J, Thiele I, Tuohy K. 2018. Gut microbiota functions: metabolism of nutrients and other food components. Eur J Nutr 57:1–24.

5. Ponomarova O, Patil KR. 2015. Metabolic interactions in microbial communities: untangling the Gordian knot. Curr Opin Microbiol 27:37–44.

6. Putnam EE, Goodman AL. 2020. B vitamin acquisition by gut commensal bacteria. PLoS Pathog 16:e1008208.

7. Warren MJ, Raux E, Schubert HL, Escalante-Semerena JC. 2002. The biosynthesis of adenosylcobalamin (vitamin B12). Nat Prod Rep 19:390–412.

8. Sokolovskaya OM, Shelton AN, Taga ME. 2020. Sharing vitamins: Cobamides unveil microbial interactions. Science 369.

9. Shelton AN, Seth EC, Mok KC, Han AW, Jackson SN, Haft DR, Taga ME. 2019. Uneven distribution of cobamide biosynthesis and dependence in bacteria predicted by comparative genomics. ISME J 13:789–804.

10. Degnan Patrick H. Taga Michiko E. Goodman AL. 2014. Vitamin B12 as a modulator of gut microbial ecology. Cell Metab 20:1–769–778.

11. Degnan PH, Barry NA, Mok KC, Taga ME, Goodman AL. 2014. Human gut microbes use multiple transporters to distinguish vitamin B□□ analogs and compete in the gut. Cell Host Microbe 15:47–57.

12. Croft MT, Lawrence AD, Raux-Deery E, Warren MJ, Smith AG. 2005. Algae acquire vitamin B12 through a symbiotic relationship with bacteria. Nature 438:90–93.

13. Doxey AC, Kurtz DA, Lynch MDJ, Sauder LA, Neufeld JD. 2015. Aquatic metagenomes implicate Thaumarchaeota in global cobalamin production. ISME J 9:461–471.

14. Cooper MB, Kazamia E, Helliwell KE, Kudahl UJ, Sayer A, Wheeler GL, Smith AG. 2019. Cross-exchange of B-vitamins underpins a mutualistic interaction between Ostreococcus tauri and Dinoroseobacter shibae. ISME J 13:334–345.

15. Lu X, Heal KR, Ingalls AE, Doxey AC, Neufeld JD. 2020. Metagenomic and chemical characterization of soil cobalamin production. ISME J 14:53–66.

16. Wang J, Shi K, Jing Z, Ge Y. 2023. Metagenomic Evidence for Cobamide Producers Driving Prokaryotic Co-occurrence Associations and Potential Function in Wastewater Treatment Plants. Environ Sci Technol 57:10640–10651.

17. Swaney MH, Sandstrom S, Kalan LR. 2022. Cobamide Sharing Is Predicted in the Human Skin Microbiome. mSystems 7:e0067722.

18. Culp EJ, Goodman AL. 2023. Cross-feeding in the gut microbiome: Ecology and mechanisms. Cell Host Microbe 31:485–499.

19. Xu Y, Xiang S, Ye K, Zheng Y, Feng X, Zhu X, Chen J, Chen Y. 2018. Cobalamin (Vitamin B12) Induced a Shift in Microbial Composition and Metabolic Activity in an in vitro Colon Simulation. Front Microbiol 9:2780.

20. Kelly CJ, Alexeev EE, Farb L, Vickery TW, Zheng L, Eric L C, Kitzenberg DA, Battista KD, Kominsky DJ, Robertson CE, Frank DN, Stabler SP, Colgan SP. 2019. Oral vitamin B12 supplement is delivered to the distal gut, altering the corrinoid profile and selectively depleting Bacteroides in C57BL/6 mice. Gut Microbes 10:654–662.

21. Zhu X, Xiang S, Feng X, Wang H, Tian S, Xu Y, Shi L, Yang L, Li M, Shen Y, Chen J, Chen Y, Han J. 2019. Impact of Cyanocobalamin and Methylcobalamin on Inflammatory Bowel Disease and the Intestinal Microbiota Composition. J Agric Food Chem 67:916–926.

22. Swaney MH, Nelsen A, Sandstrom S, Kalan LR. 2023. Sweat and Sebum Preferences of the Human Skin Microbiota. Microbiol Spectr 11:e0418022.

23. Salamzade R, Cheong JZA, Sandstrom S, Swaney MH, Stubbendieck RM, Starr NL, Currie CR, Singh AM, Kalan LR. 2023. Evolutionary investigations of the biosynthetic diversity in the skin microbiome using lsaBGC. Microb Genom 9.

24. Schwengers O, Jelonek L, Dieckmann MA, Beyvers S, Blom J, Goesmann A. 2021. Bakta: rapid and standardized annotation of bacterial genomes via alignment-free sequence identification. Microb Genom 7.

25. Aramaki T, Blanc-Mathieu R, Endo H, Ohkubo K, Kanehisa M, Goto S, Ogata H. 2020. KofamKOALA: KEGG Ortholog assignment based on profile HMM and adaptive score threshold. Bioinformatics 36:2251–2252.

26. Küberl A, Mengus-Kaya A, Polen T, Bott M. 2020. The Iron Deficiency Response of Corynebacterium glutamicum and a Link to Thiamine Biosynthesis. Appl Environ Microbiol 86.

27. Datsenko KA, Wanner BL. 2000. One-step inactivation of chromosomal genes in Escherichia coli K-12 using PCR products. Proc Natl Acad Sci U S A 97:6640–6645.

28. Chen S, Zhou Y, Chen Y, Gu J. 2018. fastp: an ultra-fast all-in-one FASTQ preprocessor. Bioinformatics 34:i884–i890.

29. Garrison E, Marth G. 2012. Haplotype-based variant detection from short-read sequencing. arXiv [q-bioGN].

30. Kelley LA, Mezulis S, Yates CM, Wass MN, Sternberg MJE. 2015. The Phyre2 web portal for protein modeling, prediction and analysis. Nat Protoc 10:845–858.

31. Brown NP, Leroy C, Sander C. 1998. MView: a web-compatible database search or multiple alignment viewer. Bioinformatics 14:380–381.

32. Jumper J, Evans R, Pritzel A, Green T, Figurnov M, Ronneberger O, Tunyasuvunakool K, Bates R, Žídek A, Potapenko A, Bridgland A, Meyer C, Kohl SAA, Ballard AJ, Cowie A, Romera-Paredes B, Nikolov S, Jain R, Adler J, Back T, Petersen S, Reiman D, Clancy E, Zielinski M, Steinegger M, Pacholska M, Berghammer T, Bodenstein S, Silver D, Vinyals O, Senior AW, Kavukcuoglu K, Kohli P, Hassabis D. 2021. Highly accurate protein structure prediction with AlphaFold. Nature 596:583–589.

33. Ittisoponpisan S, Islam SA, Khanna T, Alhuzimi E, David A, Sternberg MJE. 2019. Can Predicted Protein 3D Structures Provide Reliable Insights into whether Missense Variants Are Disease Associated? J Mol Biol 431:2197–2212.

34. Rodionov DA, Arzamasov AA, Khoroshkin MS, Iablokov SN, Leyn SA, Peterson SN, Novichkov PS, Osterman AL. 2019. Micronutrient Requirements and Sharing Capabilities of the Human Gut Microbiome. Front Microbiol 10:1316.

35. Garst AD, Edwards AL, Batey RT. 2011. Riboswitches: Structures and mechanisms. Cold Spring Harbor Perspectives in Biology. Cold Spring Harbor Laboratory Press 10.1101/cshperspect.a003533.

36. Nahvi A, Barrick JE, Breaker RR. 2004. Coenzyme B12 riboswitches are widespread genetic control elements in prokaryotes. Nucleic Acids Res 32:143–150.

37. Polaski JT, Webster SM, Johnson JE, Batey RT. 2017. Cobalamin riboswitches exhibit a broad range of ability to discriminate between methylcobalamin and adenosylcobalamin. J Biol Chem 292:11650–11658.

38. Cheng J, Poduska B, Morton RA, Finan TM. 2011. An ABC-type cobalt transport system is essential for growth of Sinorhizobium meliloti at trace metal concentrations. J Bacteriol 193:4405–4416.

39. Gu S, Sushko O, Deery E, Warren MJ, Pickersgill RW. 2015. Crystal structure of CobK reveals strand-swapping between Rossmann-fold domains and molecular basis of the reduced precorrin product trap. Sci Rep 5:16943.

40. Bauer CB, Fonseca MV, Holden HM, Thoden JB, Thompson TB, Escalante-Semerena JC, Rayment I. 2001. Three-dimensional structure of ATP:corrinoid adenosyltransferase from Salmonella typhimurium in its free state, complexed with MgATP, or complexed with hydroxycobalamin and MgATP. Biochemistry 40:361–374.

41. Saraste M, Sibbald PR, Wittinghofer A. 1990. The P-loop--a common motif in ATP- and GTP-binding proteins. Trends Biochem Sci 15:430–434.

42. Magnúsdóttir S, Ravcheev D, de Crécy-Lagard V, Thiele I. 2015. Systematic genome assessment of B-vitamin biosynthesis suggests co-operation among gut microbes. Front Genet 6:148.

43. Grant MAA, Kazamia E, Cicuta P, Smith AG. 2014. Direct exchange of vitamin B12 is demonstrated by modelling the growth dynamics of algal-bacterial cocultures. ISME J 8:1418–1427.

44. Bunbury F, Deery E, Sayer AP, Bhardwaj V, Harrison EL, Warren MJ, Smith AG. 2022. Exploring the onset of B12 -based mutualisms using a recently evolved Chlamydomonas auxotroph and B12 -producing bacteria. Environ Microbiol 24:3134–3147.

45. Kazamia E, Czesnick H, Van Nguyen TT, Croft MT, Sherwood E, Sasso S, Hodson SJ, Warren MJ, Smith AG. 2012. Mutualistic interactions between vitamin B12 -dependent algae and heterotrophic bacteria exhibit regulation. Environ Microbiol 14:1466–1476.

46. Helliwell KE, Collins S, Kazamia E, Purton S, Wheeler GL, Smith AG. 2015. Fundamental shift in vitamin B12 eco-physiology of a model alga demonstrated by experimental evolution. ISME J 9:1446–1455.

47. Yan J, Ritalahti KM, Wagner DD, Löffler FE. 2012. Unexpected specificity of interspecies cobamide transfer from Geobacter spp. to organohalide-respiring Dehalococcoides mccartyi strains. Appl Environ Microbiol 78:6630–6636.

48. Yan J, Im J, Yang Y, Löffler FE. 2013. Guided cobalamin biosynthesis supports Dehalococcoides mccartyi reductive dechlorination activity. Philos Trans R Soc Lond B Biol Sci 368:20120320.

49. Men Y, Seth EC, Yi S, Allen RH, Taga ME, Alvarez-Cohen L. 2014. Sustainable growth of Dehalococcoides mccartyi 195 by corrinoid salvaging and remodeling in defined lactate-fermenting consortia. Appl Environ Microbiol 80:2133–2141.

50. Ma AT, Beld J, Brahamsha B. 2017. An Amoebal Grazer of Cyanobacteria Requires Cobalamin Produced by Heterotrophic Bacteria. Appl Environ Microbiol 83.

51. Ma AT, Tyrell B, Beld J. 2020. Specificity of cobamide remodeling, uptake and utilization in Vibrio cholerae. Mol Microbiol 113:89–102.

52. Gude Sebastian, Pherribo Gordon J., Taga Michiko E. 2022. A Salvaging Strategy Enables Stable Metabolite Provisioning among Free-Living Bacteria. mSystems 7:e00288–22.

53. Smakman F, Hall AR. 2022. Exposure to lysed bacteria can promote or inhibit growth of neighboring live bacteria depending on local abiotic conditions. FEMS Microbiol Ecol 98.

54. Pherribo GJ, Taga ME. 2023. Bacteriophage-mediated lysis supports robust growth of amino acid auxotrophs. ISME J 17:1785–1788.

55. Shearer N, Hinsley AP, Van Spanning RJ, Spiro S. 1999. Anaerobic growth of Paracoccus denitrificans requires cobalamin: characterization of cobK and cobJ genes. J Bacteriol 181:6907–6913.

56. Blanche F, Thibaut D, Famechon A, Debussche L, Cameron B, Crouzet J. 1992. Precorrin-6x reductase from Pseudomonas denitrificans: purification and characterization of the enzyme and identification of the structural gene. J Bacteriol 174:1036–1042.

57. Kim W, Major TA, Whitman WB. 2005. Role of the precorrin 6-X reductase gene in cobamide biosynthesis in Methanococcus maripaludis. Archaea 1:375–384.

58. Kipkorir T, Mashabela GT, de Wet TJ, Koch A, Dawes SS, Wiesner L, Mizrahi V, Warner DF. 2021. De Novo Cobalamin Biosynthesis, Transport, and Assimilation and Cobalamin-Mediated Regulation of Methionine Biosynthesis in Mycobacterium smegmatis. J Bacteriol 203.

59. Medina C, Crespo-Rivas JC, Moreno J, Espuny MR, Cubo MT. 2009. Mutation in the cobO gene generates auxotrophy for cobalamin and methionine and impairs the symbiotic properties of Sinorhizobium fredii HH103 with soybean and other legumes. Arch Microbiol 191:11–21.

60. Johnson CL, Pechonick E, Park SD, Havemann GD, Leal NA, Bobik TA. 2001. Functional genomic, biochemical, and genetic characterization of the Salmonella pduO gene, an ATP:cob(I)alamin adenosyltransferase gene. J Bacteriol 183:1577–1584.

61. Gonzalez JC, Banerjee RV, Huang S, Sumner JS. 1992. Comparison of cobalamin-independent and cobalamin-dependent methionine synthases from Escherichia coli: two solutions to the same chemical problem. Biochemistry.

62. Xie B, Bishop S, Stessman D, Wright D, Spalding MH, Halverson LJ. 2013. Chlamydomonas reinhardtii thermal tolerance enhancement mediated by a mutualistic interaction with vitamin B12-producing bacteria. ISME J 7:1544–1555.

63. Mars Brisbin M, Schofield A, McIlvin MR, Krinos AI, Alexander H, Saito MA. 2023. Vitamin B12 conveys a protective advantage to phycosphere-associated bacteria at high temperatures. ISME Commun 3:88.

64. Bertrand EM, Moran DM, McIlvin MR, Hoffman JM, Allen AE, Saito MA. 2013. Methionine synthase interreplacement in diatom cultures and communities: Implications for the persistence of B12 use by eukaryotic phytoplankton. Limnol Oceanogr 58:1431–1450.

65. Dolan SK, Wijaya A, Geddis SM, Spring DR, Silva-Rocha R, Welch M. 2018. Loving the poison: the methylcitrate cycle and bacterial pathogenesis. Microbiology 164:251–259.

66. Lee W, VanderVen BC, Fahey RJ. 2013. Intracellular Mycobacterium tuberculosis exploits host-derived fatty acids to limit metabolic stress. Journal of Biological.

67. Bomar L, Brugger SD, Yost BH, Davies SS, Lemon KP. 2016. Corynebacterium accolens Releases Antipneumococcal Free Fatty Acids from Human Nostril and Skin Surface Triacylglycerols. MBio 7:e01725–15.

68. Ridaura VK, Bouladoux N, Claesen J, Chen YE, Byrd AL, Constantinides MG, Merrill ED, Tamoutounour S, Fischbach MA, Belkaid Y. 2018. Contextual control of skin immunity and inflammation by Corynebacterium. J Exp Med 215:785–799.

69. Ahmed Nashwa, Joglekar Payal, Deming Clayton, Lemon Katherine P., Kong Heidi H., Segre Julie A., Conlan Sean. 2023. Genomic characterization of the C. tuberculostearicum species complex, a prominent member of the human skin microbiome. mSystems 0:e00632–23.

70. Salamzade R, Swaney MH, Kalan LR. 2023. Comparative Genomic and Metagenomic Investigations of the Corynebacterium tuberculostearicum Species Complex Reveals Potential Mechanisms Underlying Associations To Skin Health and Disease. Microbiol Spectr 11:e0357822.

