## Supplemental Figures and Tables for "Skin-associated *Corynebacterium amycolatum* shares cobamides"

| Supplemental Table 1. Plasmids used in this study. | | |
|---|---|---|
| **Plasmid** | **Description** | **Reference** |
| pKD4 | *bla,* FRT-*kan*-FRT | (27) |
| pKD46 | *bla*, *repA101*(ts), *araBp-gam-bet-exo*, *oriR101*, *araC*, *[tL3]* | (27) |

| Supplemental Table 2. Primers used in this study. | | |
|---|---|---|
| **Primer Name** | **Primer Sequence (5’ -> 3’)** | **Reference** |
| MetH_Upstream_Fwd | AGTATGGTTTGTTGAATTTTTATTAAATCTGGGTTGAGCGTGTCGGGAGCGTGTAGGCTGGAGCTGCTTC | This study |
| MetH_Downstream_Rev | GAGACAGGGCACTCATTGAGCGCCCTGTTTGTTGTAAGGAAAAGTGACAGATGGGAATTAGCCATGGTCC | This study |
| k1 | CAGTCATAGCCGAATAGCCT | (27) |
| k2 | CGGTGCCCTGAATGAACTGC | (27) |
| kt | CGGCCACAGTCGATGAATCC | (27) |
| MetH_Outside_Fwd | CCATGACAGCCTCCTTTTTC | This study |
| MetH_Outside_Rev | GGCATCTTTATTGTGCCTTTTGAC | This study |

| Supplemental Table 3. Cobamide concentration comparison between indicator assay and mass spectrometry. | | | |
|---|---|---|---|
| | | *E. coli indicator assay* | *Mass spectrometry* |
| | **Strain** | **Cobamide concentration (μg/g)** | **Cyanocobalamin concentration (μg/g)** |
| *Cell extract* | *C. amycolatum* WT (0 nM $CoCl_2$) | 0.079 ± 0.003 | 0.017 ± 0.005 |
| | *C. amycolatum* WT (250 nM $CoCl_2$) | 4.58 ± 1.5 | 1.17 ± 0.37 |
| | *C. amycolatum* $cob^-$ (50 nM $CoCl_2$) | 0 | <LOQ |
| | *C. glucuronolyticum* | 0.051 ± 0.004 | <LOQ |

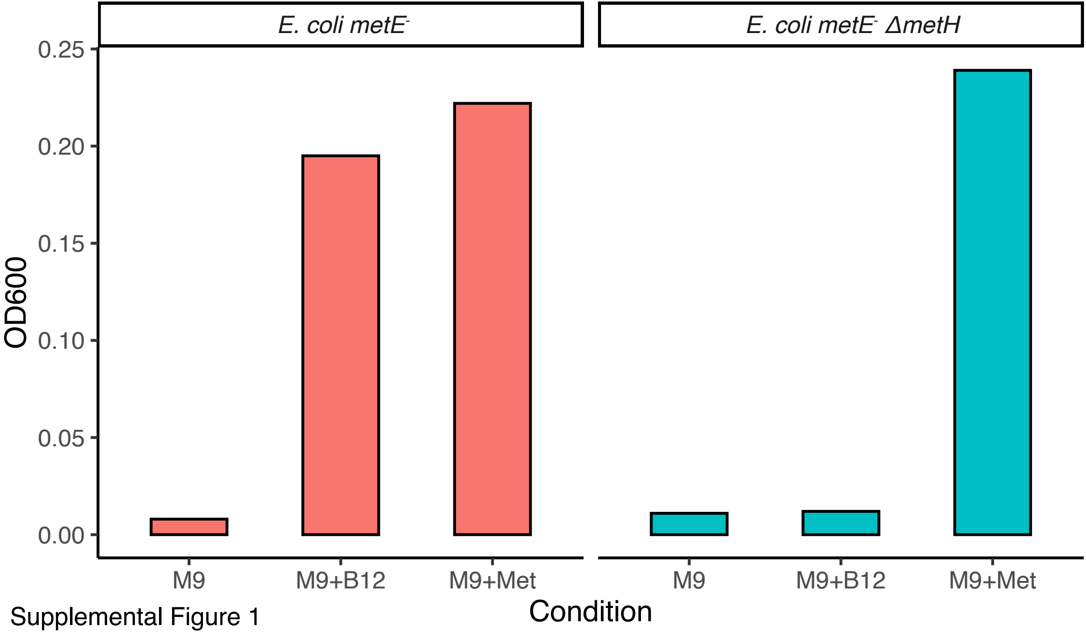


Supplemental Figure 1

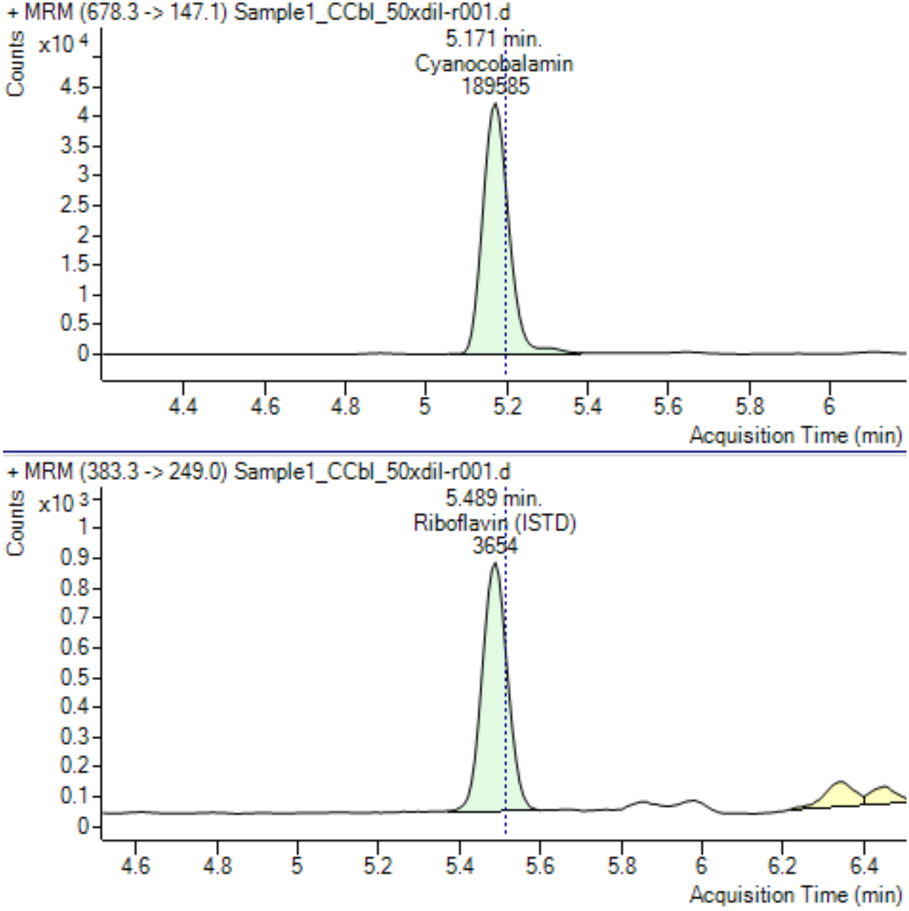


Supplemental Figure 2

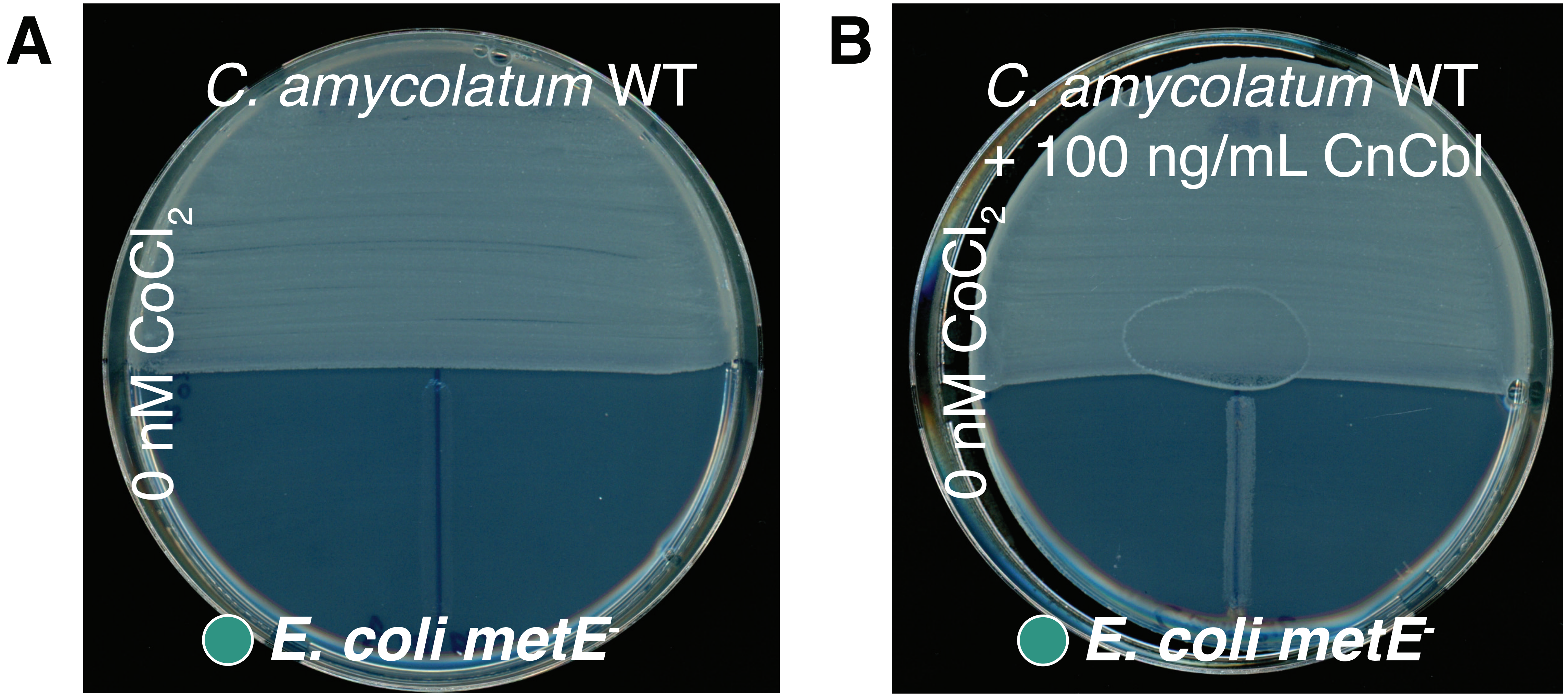


F

| Condition | 0 nM $CoCl_2$ | 250 nM $CoCl_2$ |
|---|---|---|
| *C. amycolatum* WT | - | ++ |
| *C. amycolatum* WT + 100 ng/mL CnCbl | +++ | ND |
| Control | - | - |
| 10 ng/mL CnCbl | + | + |
| 100 ng/mL CnCbl | +++ | +++ |

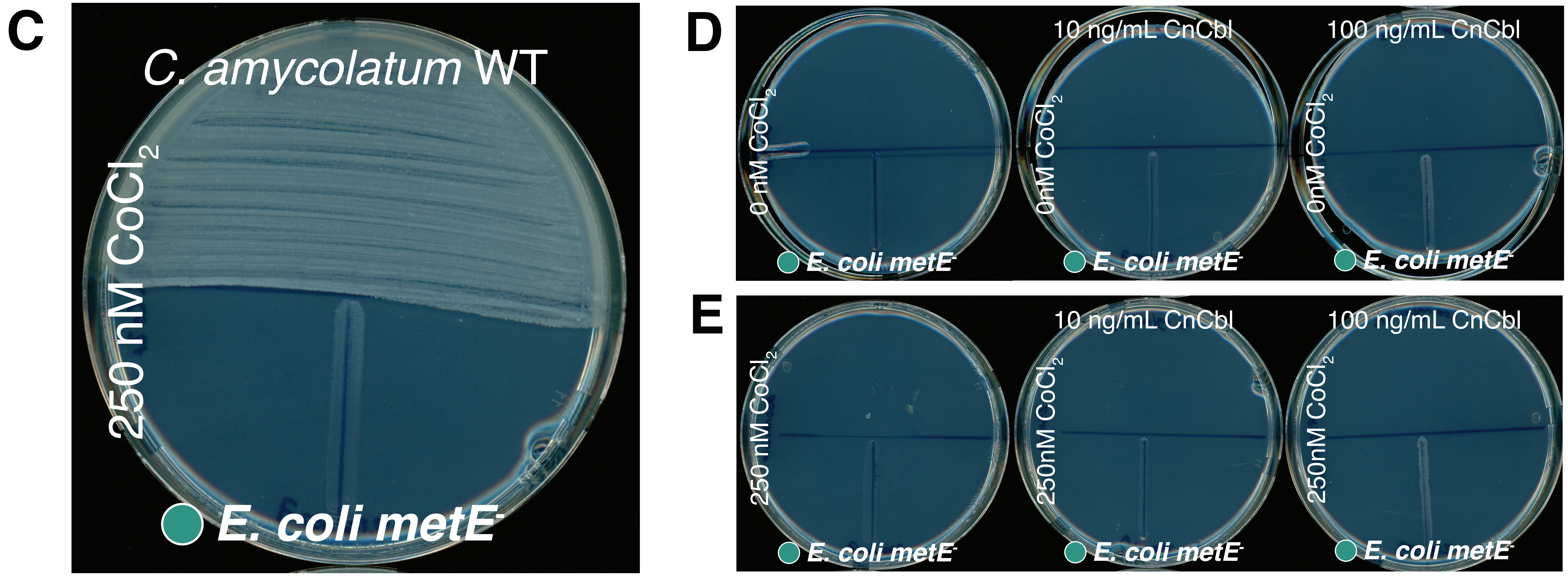


Supplemental Figure 3

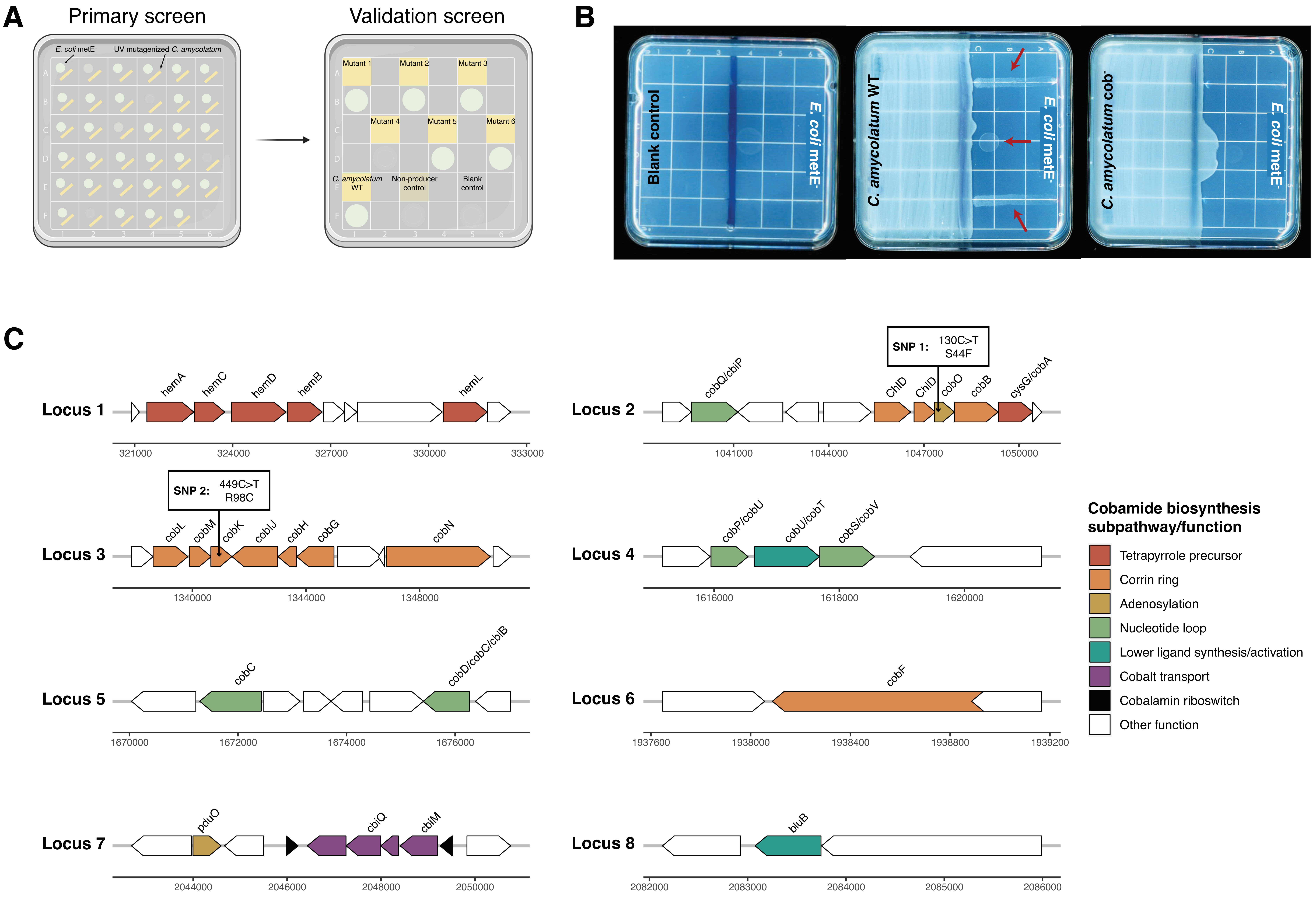


Supplemental Figure 4

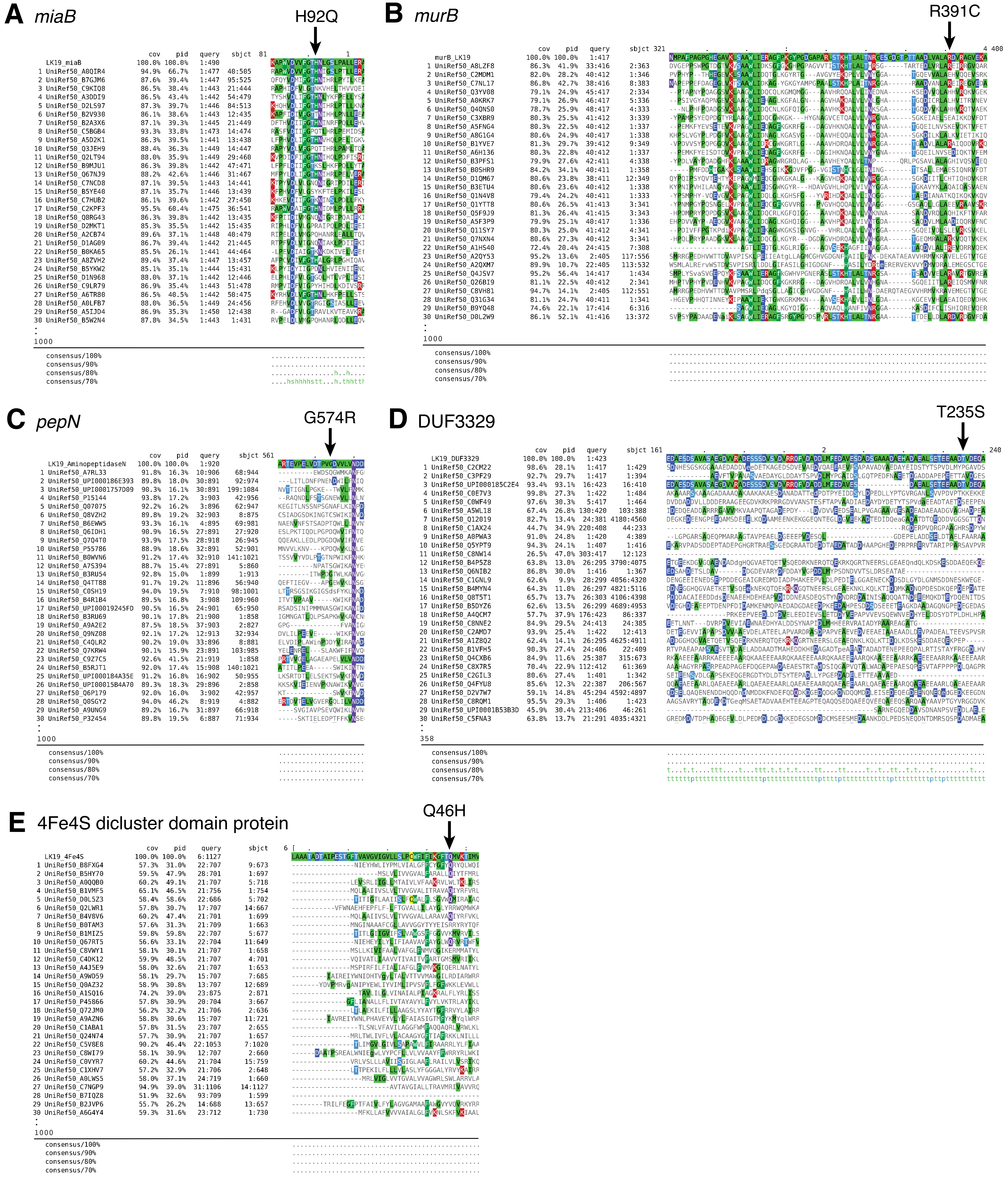


Supplemental Figure 5

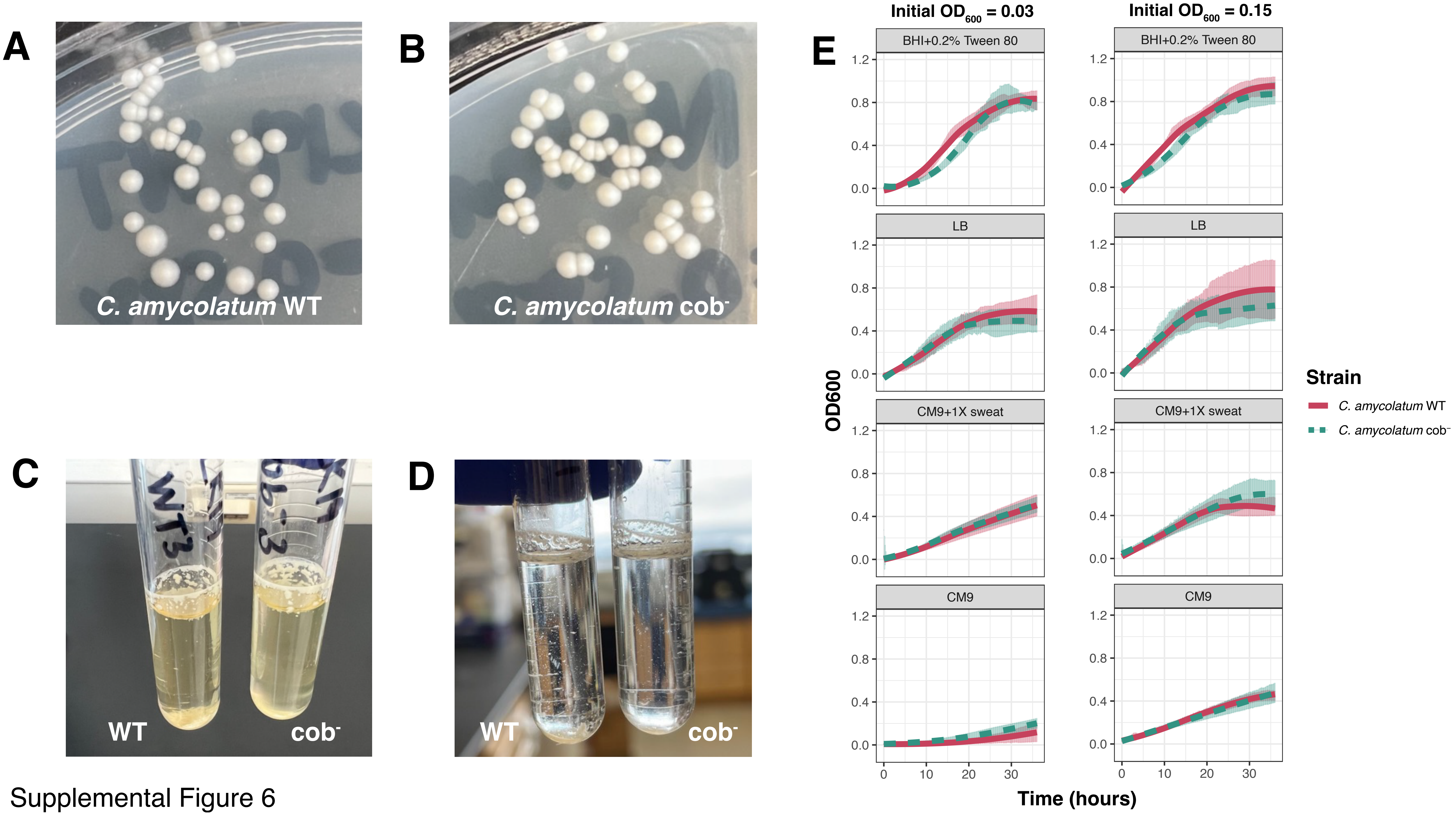


Supplemental Figure 6
